# MMPs and NETs are detrimental in human CNS-tuberculosis and MMP inhibition in a mouse model improves survival

**DOI:** 10.1101/2023.10.05.561002

**Authors:** Xuan Ying Poh, Fei Kean Loh, Chen Bai, Hai Tarng Chong, Wei Keat Teo, Jia Mei Hong, Qing Hao Miow, Pei Min Thong, Bryce Vilaysane, Ting Huey Hu, Srishti Chhabra, Yu Wang, Siew Ching Tiong, Siew Moy Fong, Masako Kamihigashi, Ravisankar Rajarethinam, Wen Donq Looi, Esther Sok Hwee Cheow, Glenn Kunnath Bonney, Leroy Sivappiragasam Pakkiri, Chester Lee Drum, Yan Fen Peng, Ming Lee, Char Loo Tan, Cristine Szu Lyn Ding, Tchoyoson Choie Cheio Lim, Tsin Wen Yeo, Joshua K Tay, Andres F. Vallejo, Catherine W M Ong

## Abstract

Despite anti-tuberculous treatment (ATT), central nervous system tuberculosis (CNS-TB) still cause permanent neurological deficits and death. To identify prognostic factors, we profiled a prospective cohort of tuberculous meningitis (TBM) and non-TBM patients. We determined significantly increased cerebrospinal fluid (CSF) matrix metalloproteinases (MMPs) and neutrophil extracellular traps (NETs) are up-regulated in TBM patients with neuroradiological abnormalities and poor outcomes. To dissect mechanisms, we created a CNS-TB murine model which show neutrophil-rich necrotizing pyogranulomas with MMP-9 and NETs colocalizing, resembling human CNS-TB. Spatial transcriptomic analysis of both human and murine CNS-TB demonstrates a highly-inflamed and neutrophil-rich microenvironment of inflammatory immune responses, extracellular matrix degradation and angiogenesis within CNS-TB granulomas. Murine CNS-TB treated with ATT and MMP inhibitors SB-3CT or doxycycline show significantly suppressed NETs with improved survival. MMP inhibition arms show attenuated inflammation and well-formed blood vessels within granulomas. Adjunctive doxycycline is highly promising to improve CNS-TB outcomes and survival.

## Introduction

Central nervous system tuberculosis (CNS-TB) is a devastating, often fatal disease that disproportionately afflicts children and immunocompromised individuals^1^. Immunopathology in CNS-TB, characterized by extensive brain inflammation and tissue destruction, is driven by a matrix-degrading phenotype, which results from an increase in matrix metalloproteinases (MMPs) relative to their tissue-specific inhibitors (tissue inhibitor of metalloproteinases, TIMPs)^2, 3^. MMPs degrade extracellular matrix (ECM) and are key to blood-brain barrier (BBB) breakdown and brain tissue damage^4^. In particular MMP-9, a gelatinase responsible for the degradation of type IV collagen, the major ECM component of the BBB, was found to be increased in the cerebrospinal fluid (CSF) of adult CNS-TB patients, which was associated with neurological deficit and death^2^. MMPs are implicated in driving CNS-TB immunopathology.

Ischemic strokes occur in up to 45% of CNS-TB patients^5^, but the mechanism of thrombosis leading to strokes in CNS-TB remains uncharacterized. Neutrophil extracellular traps (NETs), which are a meshwork of chromatin fibers studded with antimicrobial peptides and proteases, maybe implicated, linking inflammation, infection and thrombosis^4, 6^. Upon *Mycobacterium tuberculosis* (*M. tb*) infection, neutrophils produce NETs to trap mycobacteria but are unable to kill the pathogen^7^. Instead, NETs can cause local host tissue damage or thrombosis if present in the circulation. The presence of NETs in respiratory samples and necrotic lung lesions of pulmonary TB patients implicates NETs in pulmonary TB pathogenesis^8, 9^. However, the pathological role of NETs in CNS-TB remains to be explored.

To better understand the immunopathogenesis in CNS-TB, we first studied human clinical samples and then developed a murine CNS-TB model that recapitulated the brain pathology and neurological symptoms observed in humans^10^. Concurrently, we examined the role of MMPs and NETs in human CNS-TB immunopathology in driving CNS-TB disease severity. CSF MMPs and NETs in a cohort of human pediatric tuberculous meningitis (TBM) patients were analyzed together with neuroimaging and compared to clinical outcomes of severe neurological deficit and death. Spatial transcriptomic profiling elucidated the differential gene expression and pathways associated with human and murine CNS-TB immunopathology. Critically, we demonstrate the efficacy of adjunctive MMP inhibition together with anti-TB treatment (ATT) as host-directed therapy in the murine CNS-TB model.

## Results

### Up-regulated human CSF MMPs and NETs strongly associate with neuroradiological abnormalities and poor clinical outcomes in human TBM

CSF from 72 human pediatric patients with TBM and non-TBM causes of CNS diseases (Supplementary Table 1, Supplementary Table 2) were analyzed for expression of MMPs and their functional activity, NETs concentration determined and correlated with neuroimaging and clinical outcomes (Fig. 1a). None of the TBM patients had HIV nor were immunosuppressed. Consistent with previous reports^2, 11–15^, MMP-9 is higher in TBM than non-TBM by 11.3-fold (*P* = 0.0007) (Fig. 1b). MMP-2 and Extracellular MMP inducer (EMMPRIN), an activator of MMPs, are 3.5-fold (*P* = 0.0369) and 7.3-fold (*P* = 0.0182) higher in TBM than non-TBM respectively (Fig. 1b). Since net proteolysis is affected by relative local MMPs and TIMPs concentrations, we also determined the concentrations of TIMP-2 and -4, which inhibit MMP-2 and -9 respectively^16, 17^. CSF TIMP-2 (*P* = 0.0047) and TIMP-4 (*P* = 0.0338) concentrations in TBM patients are significantly down-regulated relative to non-TBM patients (Fig. 1c). This disproportionate increase in CSF gelatinases relative to TIMPs extends the previous finding in adult TBM patients^2^, and demonstrates the presence of a matrix-degrading phenotype in both pediatric and adult CNS-TB. CSF MMP-2 and -9 of TBM patients are demonstrated to be functionally active and the addition of neutralizing antibodies significantly reduce the functional gelatinase activity (MMP-2, *P* = 0.0078; MMP-9, *P* = 0.0007), confirming that the ECM degradation in the CSF is in part driven by MMP-2 and -9 (Fig. 1d).

**Fig. 1.**
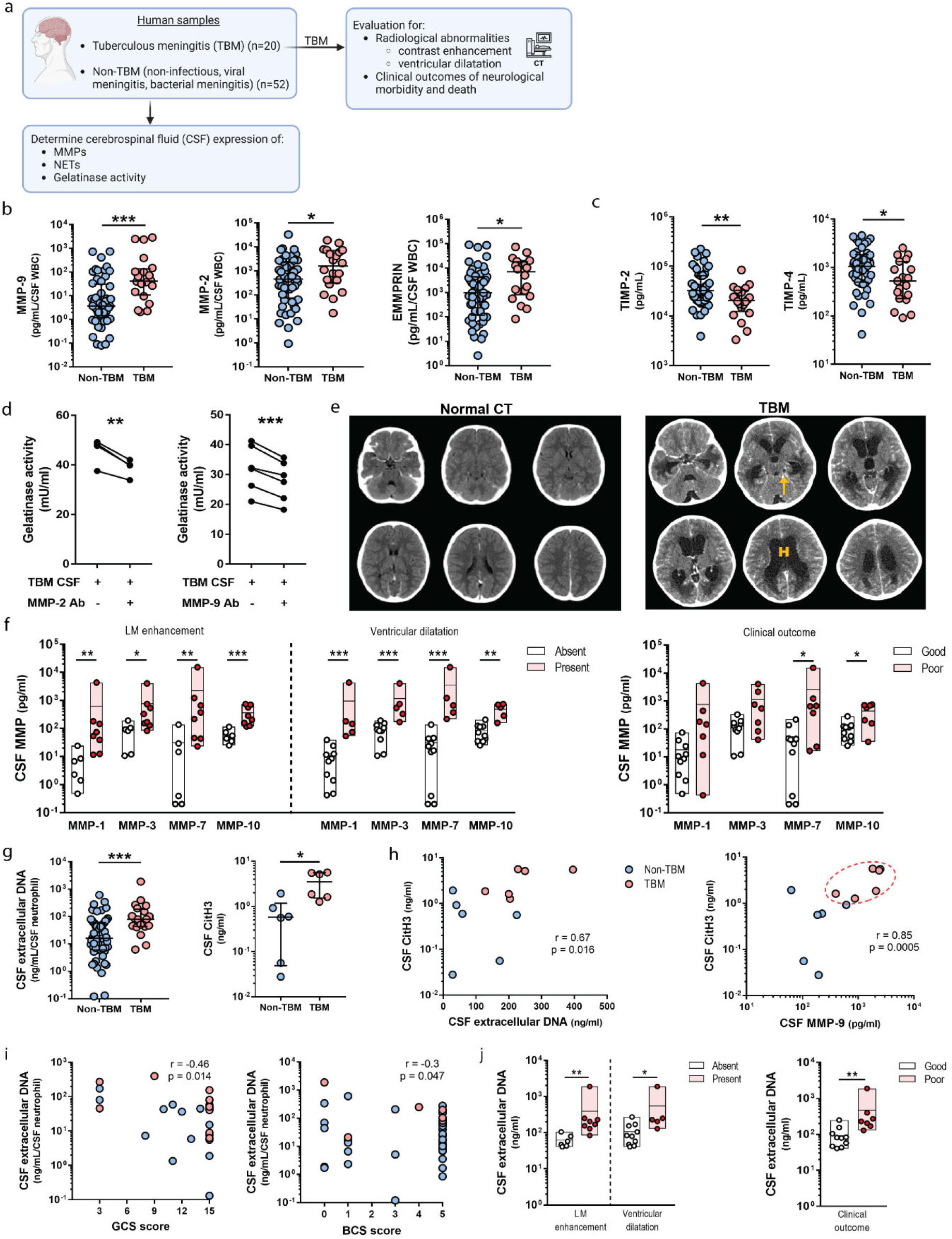
Up-regulated MMP and NET concentration in human pediatric TBM patients are strongly associated with neuroradiological abnormalities and poor clinical outcomes. **a,** CSF samples from a pediatric cohort of TBM patients and non-TBM patients were analyzed for the expression of MMPs, NETs, and functional gelatinase activity. The CT brain images of TBM patients were assessed for radiological abnormalities together with clinical outcomes. Illustration created using Biorender. **b,** CSF concentrations of MMP-9, MMP-2 and EMMPRIN are upregulated in human pediatric TBM. **c,** CSF concentration of TIMP-2 and TIMP-4 are downregulated in TBM. In **b, c**, bars represent median ± IQR with analysis by Mann-Whitney test. Non-TBM, n=52; TBM, n = 20. **d,** MMP-2 and -9 neutralization significantly suppresses CSF gelatinase activity in TBM patients. Analysis by paired t-test. MMP-2, n = 4; MMP-9, n = 6. **e,** Representative CT brain imaging of human pediatric TBM patients. Left panel shows a normal CT brain in pediatric TBM. Right panel shows widespread leptomeningeal enhancement (arrow) and dilated ventricles from hydrocephalus (H) in a pediatric TBM. **f,** Increased CSF MMPs significantly associate with leptomeningeal enhancement, ventricular dilatation and poor clinical outcomes in human TBM. Bars represent median ± IQR, dots represent individual values. Analysis by Mann-Whitney test. **g,** NETs marker extracellular DNA (n = 52 non-TBM, 20 TBM) and citH3 (n = 6 per group) are increased in human TBM. Bars represent median ± IQR. Analysis by Mann-Whitney test. **h,** CSF CitH3 concentration strongly associates with CSF extracellular DNA concentration and CSF MMP-9 concentrations. TBM patients form a cluster with high NET and MMP-9 concentrations (circled). Analysis by Pearson correlation coefficient. **i,** High CSF extracellular DNA is associated with lower GCS and BCS scores. Analysis by Pearson correlation coefficient. **j,** High CSF extracellular DNA significantly associate with leptomeningeal enhancement, ventricular dilation and poor clinical outcomes in human TBM. Bars represent median ± IQR, dots represent individual values. Statistical analysis was conducted using Mann-Whitney test. * *P* < 0.05; ** *P* < 0.01, *** *P* < 0.001.

Next, we determined the clinical significance of CSF MMP concentrations in TBM patients. Contrast enhancement, a marker of BBB disruption, and ventricular dilatation (hydrocephalus) are established poor prognostic factors,^18^ and are observed in TBM patients (Fig. 1e). Four MMPs are found to be significantly elevated in TBM patients with leptomeningeal contrast enhancement (collagenase MMP-1, *P* = 0.0027; stromelysin MMP-3, *P* = 0.0213; matrilysin MMP-7, *P* = 0.008; stromelysin MMP-10, *P* = 0.0007) and ventricular dilatation (MMP-1, *P* = 0.0007; MMP-3, *P* = 0.001; MMP-7, *P* = 0.0007; MMP-10, *P* = 0.0013) (Fig. 1f). In patients with poor clinical outcomes (survival with severe neurological deficit or death), CSF MMP-10 (*P* = 0.0243) and MMP-7 (*P* = 0.0185) concentrations are significantly higher than those with good clinical outcome (full recovery or good recovery with mild neurological deficit), while MMP-3 and -1 show a trend to increase (Fig. 1f). These findings highlight that increased CSF MMP concentrations are associated with neuroradiological abnormalities and poor clinical outcomes in TBM patients.

As NETs are implicated in TB pathogenesis, we next determined NETs in the human cohort and found extracellular DNA to be 4.9-fold higher in TBM than non-TBM patients (*P* = 0.0006) (Fig. 1g). As extracellular DNA concentration may include bacterial DNA, we further measured citrullinated histone 3 (CitH3), a NETs marker, and verified a 6-fold higher of CSF CitH3 in TBM than non-TBM patients (*P* = 0.0152) (Fig. 1g). The significant positive and strong association between CSF CitH3 and extracellular DNA concentrations (r = 0.67, *P* = 0.016) indicate that extracellular DNA concentration is a good surrogate marker for NETs (Fig. 1h). MMP-9, which degrades type IV collagen in the BBB and is established to be associated with NETs^7, 19, 20^, also demonstrates a significant positive correlation with CitH3 (r = 0.85, *P* = 0.0005), with TBM patients forming a cluster of higher NETs and MMP-9 compared to non-TBM patients (Fig. 1h).

Next, we determined whether elevated CSF NETs concentration is a poor prognostic factor. Independent of etiology, we observe a correlation of falling GCS (r = -0.46, *P* = 0.014) and BCS score (r = -0.3, *P* = 0.047) with higher CSF NETs (Fig. 1i). Interestingly, we consistently found elevated CSF extracellular DNA concentration in TBM patients with leptomeningeal enhancement (*P* = 0.0013), ventricular dilatation (*P* = 0.0193) and poor clinical outcome (*P* = 0.002) (Fig. 1j). These findings indicate that raised CSF NETs concentration is a marker of poor prognosis in TBM.

### MMP-2 and -9 are upregulated in H37Rv-infected C57BL/6 *Nos2^-/-^*mice with increased extracellular matrix degradation and NETs

We determined a suitable murine strain for CNS-TB^10^ by comparing C57BL6 *Nos2^-/-^*with the C3HeB/FeJ mice, which is known to be a good murine pulmonary TB model^21^. Mice were infected with *M. tb* H37Rv for 8 weeks (Fig. 2a). We found MMP-2 and -9 concentrations in infected *Nos2*^-/-^ mice to be 3.3-fold (*P* = 0.0075) and 40-fold (*P* = 0.011) higher than infected C3HeB/FeJ mice brain (Fig. 2b). Apart from brain gelatinases, stromelysin MMP-3 and elastase MMP-12 concentrations in *Nos2*^-/-^ mice are also significantly up-regulated compared to C3HeB/FeJ mice by 21-fold (*P* = 0.013) and 49-fold (*P* = 0.016) respectively (Fig. 2c). In C3HeB/FeJ mice, there is no difference between infected and saline controls in all MMPs analyzed (Fig. 2b, Fig 2c).

**Fig. 2.**
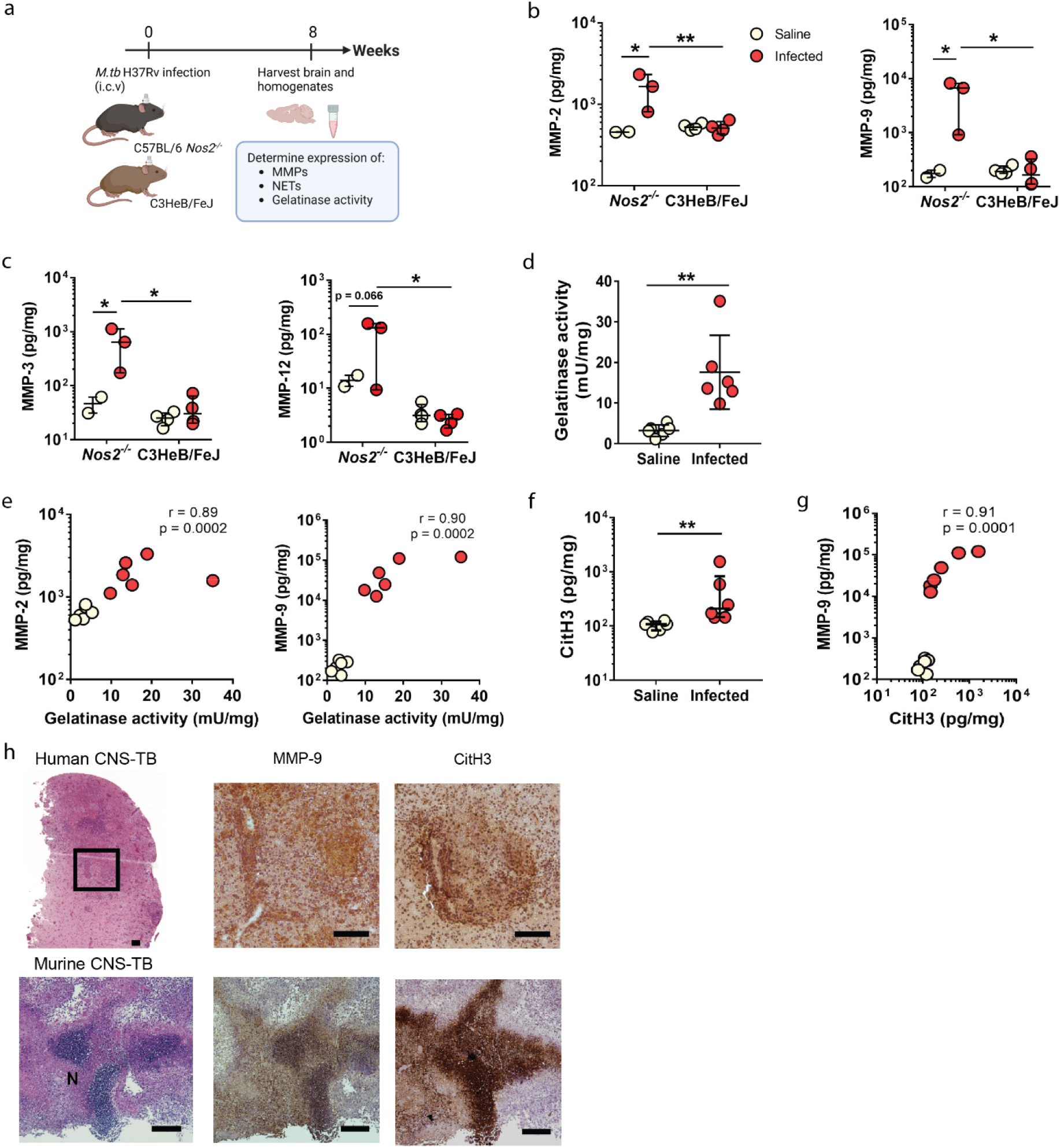
Functional gelatinase activity is increased with H37Rv infection and is associated with MMP-2 and -9 concentrations in murine CNS-TB. **a,** Six- to eight-week-old male C57BL6 *Nos2^-/-^* and C3HeB/FeJ mice were infected with 10^5^ CFU *M. tb* H37Rv via intracerebroventricular (i.c.vent). route, control mice received saline (n = 2 to 4). Eight weeks post-infection, the brain tissues and homogenates were used for downstream assay. Illustration created using Biorender. Brain concentrations of (**b**) gelatinase MMP-2, -9, and (**c**) stromelysins MMP-3, elastase MMP-12 are upregulated in H37Rv i.c.vent.-infected C57BL6 *Nos2^-/-^* compared to C3HeB/FeJ mice. MMP concentrations are normalized to total protein concentration. Bars represent median ± IQR, and analysis was conducted using two-way ANOVA with Sidak’s multiple comparisons test. **d**, Functional gelatinase activity in the brain homogenates of H37Rv i.c.vent.-infected *Nos2^-/-^* mice is significantly increased. Bars represent mean ± SD. Analysis by unpaired t test. **e**, Brain gelatinase activity are associated with MMP-2 and MMP-9 concentrations, analyzed by Spearman’s correlation coefficient. **f**, Brain concentration of NETs marker CitH3 is significantly up-regulated in H37Rv i.c.vent.-infected *Nos2^-/-^* mice relative to saline control. Bars represent median ± IQR. Analysis by Mann-Whitney test. **g,** Brain MMP-9 is associated with NETs marker CitH3 concentrations. Analysis by Spearman’s correlation coefficient. * *P* < 0.05; ** *P* < 0.01. **h,** IHC stains of granulomas in human CNS-TB and murine CNS-TB show expression of MMP-9 and CitH3 at the necrotic (N) area. Top panel, scale bar = 100 µm; Bottom panel, scale bar = 200 µm.

We replicated the study with a larger sample size (n = 6) and similarly show significantly increased MMP-2 (2.8-fold, *P* = 0.0022), MMP-9 (154.2-fold, *P* = 0.0022), MMP-3 (347.3-fold, *P* = 0.0022) and MMP-12 (122.3-fold, *P* = 0.0022) in the *M. tb*-infected C57BL/6 *Nos2^-/-^* mice relative to saline control (Supplementary Fig. 1a). In H37Rv-infected mice, TIMP-2 concentration is increased relative to saline control (*P* = 0.0411), while TIMP-4 concentration tends to decrease relative to saline control, but do not reach significant difference (*P* = 0.1797) (Supplementary Fig. 1b). Overall, these findings support the use of C57BL/6 *Nos2^-/-^* mice as a murine CNS-TB model with a destructive matrix-degrading phenotype that reflects human disease.

While TIMPs are widely recognized as the endogenous inhibitors of MMPs, TIMP-2 is also involved in MMP-2 activation^22, 23^. Thus, we sought to determine the net proteolytic activity in the murine brain by determining the functional gelatin degradation. Brain gelatinase activity is significantly increased in H37Rv-infected *Nos2^-/-^* mice relative to saline control by 5.5-fold (*P* = 0.0033) (Fig. 2d). Gelatinase activity significantly associate with MMP-2 (r = 0.8932, *P* = 0.0002) and MMP-9 (r = 0.9002, *P* = 0.0002) concentrations (Fig. 2e). This indicates that H37Rv infection in murine CNS-TB results in the development of a tissue destructive microenvironment.

To determine if our murine model has further similarities to human CNS-TB, we measured NETs in brain homogenates. CitH3 is significantly upregulated in H37Rv-infected *Nos2^-/-^* mice compared to saline control by 1.9-fold (*P* = 0.0022) (Fig. 2f). Similar to human TBM, MMP-9, demonstrates a significant association with CitH3 (r = 0.91, *P* = 0.0001) (Fig. 2g). Immunohistochemistry confirms high expression of MMP-9 and CitH3 in the necrotic region of tuberculomas in both human and murine CNS-TB (Fig. 2h). Together, the upregulated MMPs and NETs during infection further supports the use of the *Nos2^-/-^*strain as our murine CNS-TB model.

### Spatial transcriptomics analysis reveals human and murine CNS-TB granuloma microenvironment is highly inflamed with elevated immune, cytokine and defense responses

To further determine underlying drivers of immunopathology within the tissue microenvironment, we performed laser capture microdissection RNA sequencing (LCM RNA-seq) to profile the transcriptome of granulomas and inflammatory regions compared to respective controls in human and murine CNS-TB (Fig. 3a). Human demyelination tissues comprising inflammatory lesions from damaged myelin sheaths^24^ (Supplementary Fig. 2a) were used as control for human samples. Normal tissues from healthy C57BL/6 *Nos2^-/-^*mice were used as control for murine samples. Hierarchical clustering performed using the top 500 variable genes across the human samples distinguished the gene expression of human CNS-TB granulomas from human demyelination controls, normal-adjacent tissues from human CNS-TB granulomas and CNS-TB inflammatory tissues (Fig. 3b, Supplementary File 1). In human CNS-TB granulomas, gene clusters show enrichment of biological processes related to epithelial mesenchymal transition, TNFα signaling via NFκB, angiogenesis, apoptosis and IFN-ɣ response (Supplementary Fig. 2b). To determine disease-specific genes and pathways in CNS-TB, we identified the differential expressed genes (DEGs) in human CNS-TB granulomas compared to control human demyelination tissues. Principal component analysis shows that both groups had distinct gene expression profiles (Supplementary Fig. 2c). A total of 108 genes are differentially expressed (*FDR* < 0.05) (Fig. 3c, Supplementary File 2). Processes enriched in human CNS-TB granulomas include cytokine responses, cell-substrate adhesion, ECM organization, inflammatory, immune responses, angiogenesis and TNFα signaling via NFκB (Supplementary Fig. 2d, Supplementary Fig. 2e). On the other hand, CNS-related processes are disenriched including synapse and cell junction organization, myelination, neuron development and differentiation (Supplementary Fig. 2f).

**Fig. 3.**
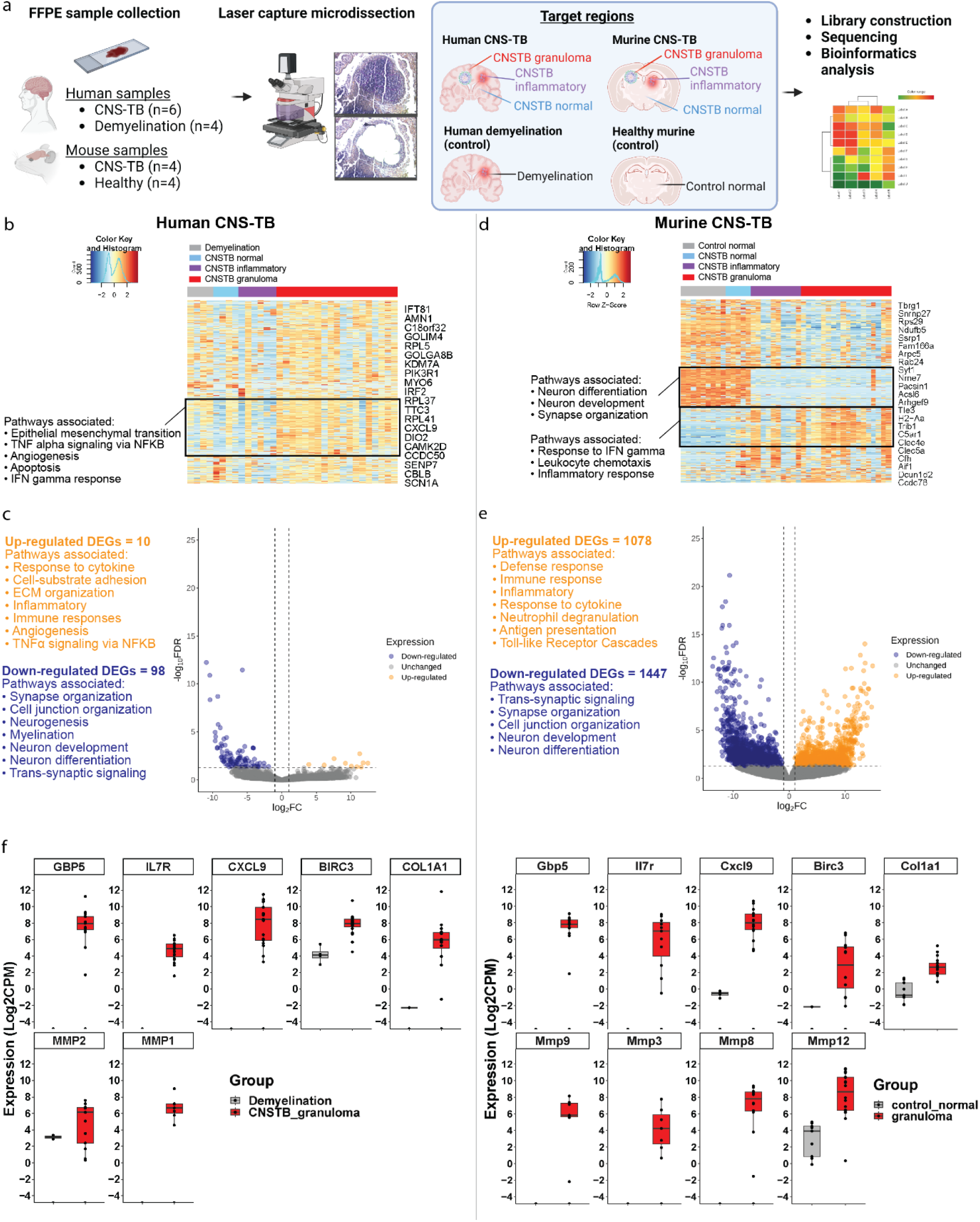
Spatial transcriptomic profiling of human and murine CNS-TB shows enrichment of inflammatory and immune genes and pathways. **a**, Target regions of formalin-fixed paraffin embedded (FFPE) CNS tissue samples from human CNS-TB (n = 6), control human demyelination tissues (n = 4), murine CNS-TB (n=4) and healthy murine controls (n = 4) were microdissected for LCM RNA-Seq. Illustration created using Biorender. **b,** Hierarchical clustering of top 500 variable genes expressed in human CNS-TB. Black frame highlights gene cluster that was distinctly expressed in human CNS-TB granulomas compared to other tissues. **c,** Volcano plot of up-regulated (log fold-change > 2, *FDR* < 0.05) and down-regulated (log fold-change < -2, *FDR* < 0.05) genes in human CNS-TB granulomas compared to human demyelination tissues. **d,** Hierarchical clustering of top 500 variable genes expressed in murine samples. Black frame highlights gene clusters that were distinctly expressed in murine CNS-TB granulomas relative to other tissue types. **e,** Volcano plot of up-regulated (log fold-change > 2, *FDR* < 0.05) and down-regulated (log fold-change < 2, *FDR* < 0.05) genes in murine CNS-TB granulomas compared to normal tissues from healthy murine controls. **f,** Genes consistently up-regulated in human and murine CNS-TB granuloma (top panel), as well as selected MMP genes up-regulated in human and murine CNS-TB granulomas (bottom panel). Box represents 25th and 75th percentile, line represents median and whiskers denote the extremes. Box represents 25th and 75th percentile, line represents median and whiskers denote the extremes.

We interrogated the gene expression of murine CNS-TB in a similar fashion (Fig. 3d, Supplementary File 3). Hierarchical clustering of the top 500 variable genes across murine CNS-TB samples shows consistent downregulation of genes associated with neuron differentiation, neuron development and synapse organization (Supplementary Fig. 3a), and enrichment of genes related to IFN-ɣ response, leukocyte chemotaxis and inflammatory response in CNS-TB granulomas and inflammatory tissues (Supplementary Fig. 3b). Principal component analysis shows distinct gene expression clusters separating murine CNS-TB granulomas and normal tissue from healthy mice (Supplementary Fig. 3c). A total of 2525 DEGs genes are differentially expressed between the two groups (*FDR* < 0.05) (Fig. 3e, Supplementary File 4). The up-regulated DEGs are implicated in defense, immune, inflammatory and cytokine responses, neutrophil degranulation, antigen presentation and Toll-Like Receptor (TLR) cascades, present in the murine CNS-TB granulomas (Supplementary Fig. 3d, Supplementary Fig. 3e). Conversely, genes related to trans-synaptic signaling, synapse and cell junction organization, neuron development and differentiation are decreased in murine CNS-TB granuloma tissues (Supplementary Fig. 3f). We observe five DEGs consistently up-regulated in human and murine CNS-TB granulomas compared to their respective controls (Fig. 3f). The genes are *GBP5*, which encodes guanylate binding protein 5, the activator of NLRP3 inflammasome^25^; *IL7R*, which encodes interleukin 7 receptor which regulates development and homeostasis of immune cells, including T cells, B cells and NK cells^26^; *CXCL9*, which encodes IFN-γ-induced chemokine which recruits T cells and NK cells^27^; *BIRC3*, which encodes Baculoviral IAP Repeat Containing 3 protein which suppresses TNF-α stimulated cell apoptosis signaling^28^; *COL1A1,* which encodes pro-alpha1 chains of type I collagen^29^. Human *MMP-2*, *-1*, and murine *Mmp-9*, *-3*, *-8* and *-12* are also up-regulated in CNS-TB granulomas, corresponding to the increased protein expression in our study (Fig. 3f).

### Neutrophil genes and associated transcriptional factors are predominantly present in human and murine CNS-TB granulomas

We sought to investigate the immune cell types in both human and murine TB granulomas. CIBERSORTx analysis was performed to infer the fractions of immune cell types in each sample^30^. Compared to human demyelination tissues, we observe significantly higher fractions of neutrophils (*P* = 0.0316), macrophages M1 (*P* = 0.0181), activated mast cells (*P* = 0.0027) and activated CD4 memory T cells (*P* = 0.0002) in human CNS-TB granulomas (Fig. 4a, Supplementary Fig. 4). The major immune cell composition in the murine CNS-TB granulomas comprises of neutrophils (*P* < 0.0001), dendritic cells (*P* = 0.0013) and NK cells (*P* < 0.0001), with higher percentage compared to normal tissues from healthy murine controls (Fig. 4b). We further examined the presence of neutrophil transcripts in murine CNS-TB granulomas^31^. Marker genes associated with nearly mature neutrophils (*Lgals3*, *Mcemp1*, *Mmp8*, *Mmp9*, *Retnlg*) and mature circulating neutrophils (*Btg1, Ccl6*, *Csf3r*, *Dusp1*, *Fth1*, *Fxyd5*, *H2-D1*, *Il1b*, *Msrb1*, *Srgn*, *Wfdc17*) are significantly up-regulated in murine CNS-TB granulomas (*FDR* < 0.05) (Fig. 4c, Supplementary File 5). These gene signatures highlight the importance of neutrophils in CNS-TB.

**Fig. 4.**
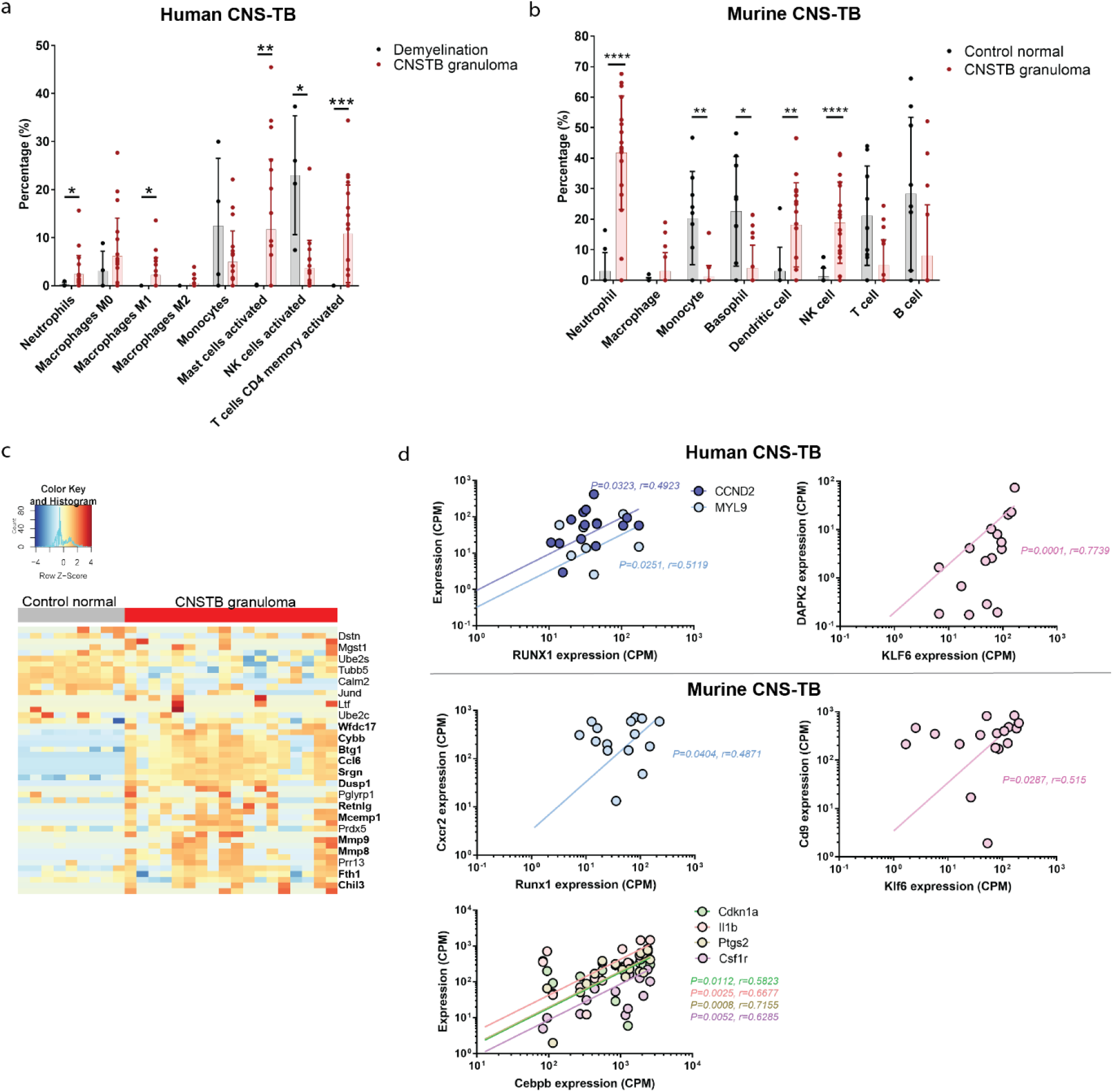
Key immune cells, neutrophil marker genes and transcription factors driving immunopathology in human and murine CNS-TB granulomas. **a**, Deconvolution analysis by CIBERSORTx LM22 signature reveals that relative to human demyelination tissue, the *in silico* cell composition in human TB granulomas has higher fractions of neutrophils, macrophages M1, activated mast cells and activated CD4 memory T cells. Bars represent mean ± SD, and analysis was conducted using unpaired t test with Welch’s correction. * *P* < 0.05; ** *P* < 0.01; *** *P* < 0.001; **** *P* < 0.0001. **b**, *In silico* deconvolution analysis using ImmuCC signature matrix indicates that relative to healthy murine brain tissues, murine CNS-TB granulomas consist mainly of neutrophils, dendritic cells and NK cells. Bars represent mean ± SD, and analysis was conducted using unpaired t test with Welch’s correction. **c**, Heatmap shows higher expression of neutrophil genes in murine CNS-TB granuloma samples compared to control normal samples. Genes in bold font are significantly upregulated with *FDR* < 0.05. **d,** Key transcriptional factors, which regulate neutrophils functions, correlate significantly with expression of their neutrophil-associated target genes in human and murine CNS-TB granulomas. Analysis by Spearman’s correlation coefficient.

Given the enrichment of neutrophils and cytokine responses in human and murine CNS-TB, we evaluated the expression of key transcription factors (TFs) known to be associated with neutrophil functions. In human CNS-TB granulomas, *RUNX1* and *KLF6*, which modulate neutrophil maturation and migration^32^, correlate significantly with neutrophil-associated target genes including *CCND2* (r = 0.4923, *P* = 0.0323), *MYL9* (r = 0.5119, *P* = 0.0251) and *DAPK2* (r = 0.7739, *P* = 0.0001) (Fig. 4d). Similarly, in murine CNS-TB granulomas, *Runx1* and *Klf6* associate significantly with their respective targets *Cxcr2* (r = 0.4871, *P* = 0.0404) and *Cd9* (r = 0.515, *P* = 0.0287) (Fig. 4d). In addition, *Cebpb*, which regulates emergency granulopoiesis and enhances *de novo* neutrophils synthesis during inflammation^33^, significantly correlates with neutrophil-associated genes *Cdkn1a* (r = 0.5823, *P* = 0.0112), *Il1b* (r = 0.6677, *P* = 0.0025), *Ptgs2* (r = 0.7155, *P* = 0.008) and *Csf1r* (r = 0.6285, *P* = 0.0052) in murine CNS-TB granulomas (Fig. 4d). These transcriptional activities implicate neutrophil-driven inflammation in human and murine CNS-TB. Together, our spatial transcriptomic analyses show marked similarities between the human and murine CNS-TB transcriptome.

### Adjunctive MMP inhibitors SB-3CT and doxycycline with standard ATT therapy significantly reduce brain immunopathology and improve murine CNS-TB survival

As both our human cohort and our CNS-TB murine model consistently demonstrate elevated MMP-2 and -9 expression, we determined if adjunctive MMP inhibitor therapy with SB-3CT or doxycycline with standard ATT therapy can suppress immunopathology in our CNS-TB murine model. The ATT treatment regime was initiated at 3 weeks post-infection for a period of 8 weeks (Fig. 5a) as the formation of brain granulomas and tissue necrosis were observed in *M. tb*-infected mice as early as 3 weeks post-infection. Compared to ATT treatment alone, both ATT with SB-3CT (*P* = 0.0135) and ATT with doxycycline (*P* = 0.0328) significantly improve survival (Fig. 5b). Brain histology shows vehicle control (infected) mice and ATT alone-treated mice contain pyogranulomas with liquefactive necrosis, while groups that received ATT with adjunctive MMP inhibitors have non-necrotizing granulomas (Fig. 5c). Furthermore, compared to vehicle control (infected) and ATT-treated groups, mice that received adjunctive SB-3CT or doxycycline demonstrate an overall reduced histopathological lesion burden (Fig. 5d, Supplementary Table 3). In groups that received ATT with MMP inhibitors, there is significant reduction of gliosis (ATT+SB-3CT, *P* = 0.0038; ATT+Doxycycline, *P* = 0.0079), pyogranulomas (ATT+SB-3CT, *P* = 0.0026; ATT+Doxycycline, *P* = 0.0134), and bacilli (ATT+SB-3CT, *P* = 0.0102; ATT+Doxycycline, *P* = 0.0102) (Fig. 5d). CFU enumeration of brain homogenates relative to ATT only group demonstrates suppressed *M. tb* growth in ATT+SB-3CT and ATT+Doxycycline group, but do not reach statistical significance (Supplementary Fig. 5). Compared to ATT only group, the total number of granulomas in ATT+SB-3CT (*P* = 0.0005) and ATT+Doxycycline groups (*P* = 0.0473) are significantly decreased (Fig. 5e). Ziehl-Neelsen (ZN) staining confirms the presence of acid-fast bacilli (AFB) in the murine CNS-TB granulomas (Fig. 5f). Notably, a smaller number of AFB positive granulomas is observed in the group that received ATT+SB-3CT relative to VC (*P* = 0.0036) or ATT alone (*P* = 0.0155), and a decreasing trend is observed in the ATT+Doxycycline group (Fig. 5g).

**Fig. 5.**
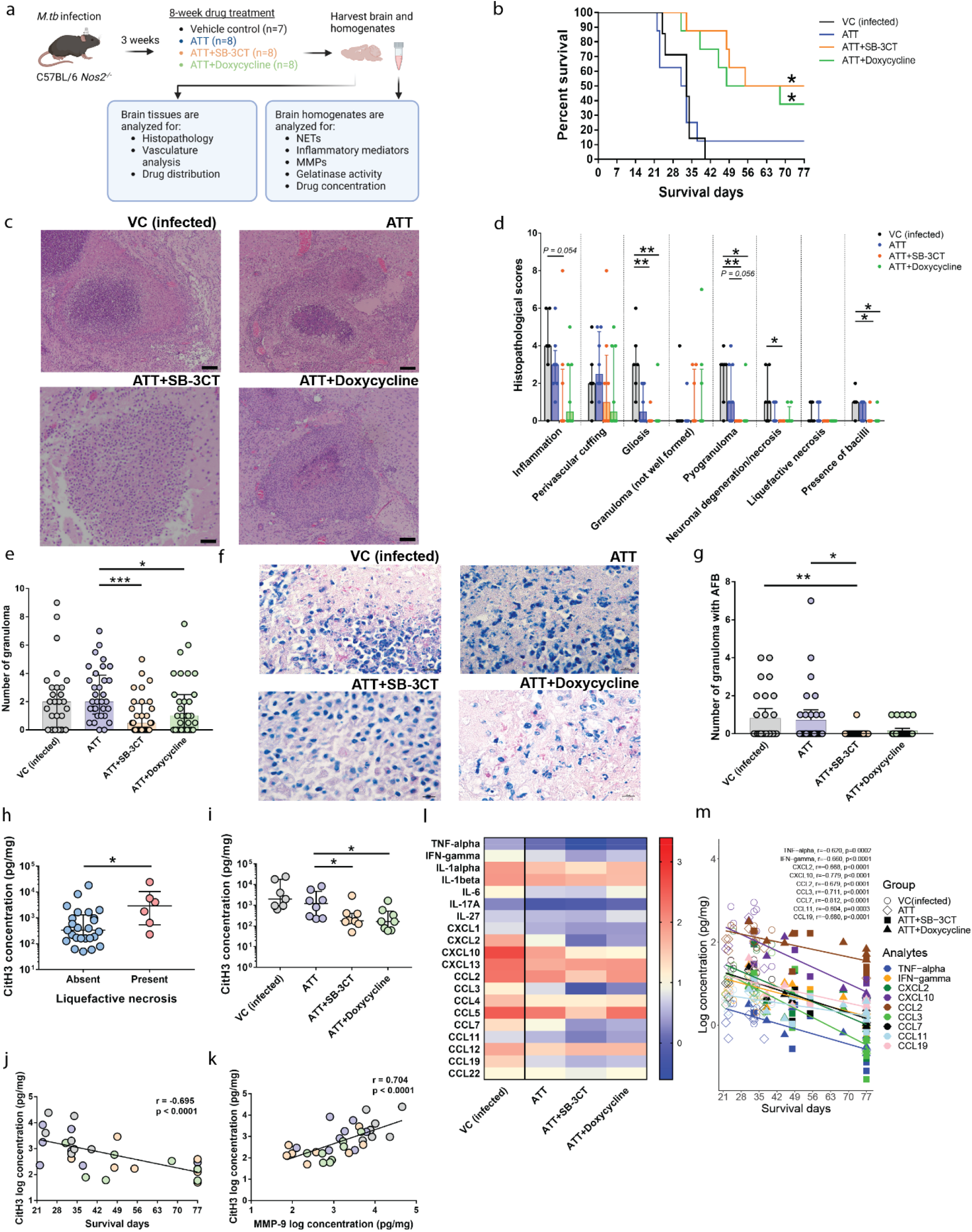
Adjunctive MMP inhibition with SB-3CT or doxycycline with standard ATT significantly improve murine CNS-TB survival and reduce immunopathology with decreased expression of NETs and inflammatory mediators. **a**, Six- to eight-week-old male C57BL/6 *Nos2^-/-^* mice were infected with 10^5^ CFU *M. tb* H37Rv via the i.c.vent. route and treated with ATT, ATT+SB-3CT or ATT+Doxycycline from 3 weeks post-infection. (n = 7 to 8 per group). After eight weeks of treatment, the brain homogenates were collected for downstream analyses. Illustration created using Biorender. **b**, Kaplan–Meier curve shows significant improved survival of *M. tb*-infected mice treated with MMP inhibitor SB-3CT or doxycycline plus ATT. **c**, Representative hematoxylin–eosin stains of *M. tb*-infected mouse brain pyogranulomas in VC (infected) and ATT alone groups (top panel), and non-necrotizing granulomas in ATT+SB-3CT and ATT+Doxycycline groups (bottom panel). Scale bar =100 µm. **d**, Histopathological severity scoring of brain lesions are higher in vehicle control (*M. tb*-infected) and ATT-alone mice with lower severity scores in adjunctive MMP inhibitor-treated mice. Bars represent median ± IQR. Analysis was conducted using Kruskal-Wallis test with Dunn’s multiple comparisons test. **e**, Granuloma numbers are less in *M. tb*-infected mice treated with ATT with adjunctive MMP inhibitors. Bars represent median ± IQR, analysis was conducted using Kruskal-Wallis test with Dunn’s multiple comparisons test. **f,** Ziehl–Neelsen staining shows presence of AFB in the granuloma of murine CNS-TB of different groups. Scale bar =10 µm. **g,** Smaller number of AFB-containing granulomas are found in mice on ATT and adjunctive MMP inhibitor SB-3CT or doxycycline. Bars represent median ± IQR., analysis was conducted using Kruskal-Wallis test with Dunn’s multiple comparisons test. **h**, CitH3 expression is significantly up-regulated in brain homogenates of mice with liquefactive necrosis. Bars represent median ± IQR. Analysis by Mann Whitney test. **i**, CitH3 expression is suppressed in CNS-TB mice treated with adjunctive MMP inhibitors SB-3CT or doxycycline. Bars represent median ± IQR. Analysis by Mann Whitney test. **j**, Decreased CitH3 in the brain homogenates of murine CNS-TB is strongly associated with longer survival days. Analysis was conducted using Spearman correlation coefficient. **k**, Brain homogenate MMP-9 is strongly associated with CitH3, analyzed by Spearman correlation coefficient. **l**, Heatmap shows suppressed median expression of inflammatory mediators in adjunctive MMP inhibitor groups. **m**, Inflammatory mediators TNF-α, IFN-ɣ, CXCL2, CXCL10, CCL2, CCL3, CCL7, CCL11, and CCL19 are inversely correlated with longer survival days (all strong associations with r < -0.5), analyzed by Spearman correlation coefficient. * *P* < 0.05; ** *P* < 0.01.

We sought to evaluate the role of NETs in our murine model and found that mice with liquefactive necrosis of brain granulomas have significantly up-regulated CitH3 (*P* = 0.0268) (Fig. 5h). After treatment with MMP inhibitors, CitH3 concentration is significantly decreased in mice treated with ATT+SB-3CT (*P* = 0.0379) and ATT+Doxycycline (*P* = 0.0207) (Fig. 5i), indicating that MMP inhibition reduces destructive NETs formation. The reduced CitH3 significantly associates with longer survival (r = -0.695, *P* < 0.0001) (Fig. 5j). We then investigated if neutrophil mediators, specifically MMP-9 and NETs, correlate to tissue destruction in CNS-TB, and found a strong association between MMP-9 and CitH3 concentrations (r = 0.704, *P* < 0.0001) (Fig. 5k).

Next, we analyzed the brain cytokines and chemokines which govern the recruitment of diverse immune cells that drive inflammation. TNF-α, a cytokine that is key to CNS-TB pathogenesis by increasing BBB permeability^34^, and other proinflammatory cytokines IFN-ɣ, IL-1α, IL-1β, IL-6, IL-17A, IL-27 are downregulated in groups that received ATT with adjunctive SB-3CT or doxycycline (Fig. 5l). Compared to vehicle control and ATT groups, neutrophil chemoattractant CXCL1 and CXCL2, monocytes and macrophages chemoattractant CCL2, CCL3, CCL7, CCL12, eosinophil chemoattractant CCL11, and Th1-associated chemokines CXCL10, CCL4, CCL5 are also down-regulated in ATT+SB-3CT and ATT+Doxycycline groups (Fig. 5l). We further demonstrate that the following suppressed cytokines and chemokines are strongly associated with longer survival; TNF-α (r = -0.620, *P* = 0.0002), IFN-ɣ (r = -0.660, *P* < 0.0001), CXCL2 (r = -0.668, *P* < 0.0001), CXCL10 ((r = - 0.779, *P* < 0.0001), CCL2 (r = -0.679, *P* < 0.0001), CCL3 (r = -0.711, *P* < 0.0001), CCL7 (r = -0.812, *P* < 0.0001), CCL11 (r = -0.604, *P* = 0.0003) and CCL19 (r = -0.680, *P* < 0.0001). Specifically, mice that received adjunctive MMP inhibitors form a cluster with lower concentrations of these inflammatory mediators relative to those without (Fig. 5m). These observations indicate that MMP inhibition suppresses pro-inflammatory cytokines and chemokines which strongly associate with improved survival outcome.

### Adjunctive MMP inhibition suppresses the matrix-degrading phenotype and normalizes vasculature in murine CNS-TB

To determine if adjunctive MMP inhibition suppresses the matrix-degrading phenotype in CNS-TB, we analyzed the brain MMPs and TIMPs of the different murine treatment groups. Relative to vehicle control and ATT-treated mice, mice that received adjunctive MMP inhibitors and ATT have decreased gelatinases MMP-2 and -9, collagenase MMP-8, stromelysin MMP-3, -10, and elastase MMP-12 (Fig. 6a). The suppressed brain MMP-3 (r = - 0.793, *P* <0.0001), MMP-8 (r = -0.662, *P* <0.0001), MMP-9 (r = -0.699, *P* <0.0001), MMP-10 (r = -0.594, *P* = 0.0004) and MMP-12 (r = -0.695, *P* <0.0001) are strongly associated with longer survival (Fig 6b). Additionally, higher MMP-3 (r = 0.457, *P* = 0.01), MMP-9 (r = 0.438, *P* = 0.014) and MMP-12 (r = 0.386, *P* = 0.032) are significantly associated with worse histopathological scores, whereas treatment groups that received adjunctive MMP inhibitors have lower histopathological scores (Fig. 6c). We find decreased TIMP-1 concentration and increased TIMP-2 and -4 concentrations in MMP inhibitors groups compared to vehicle control (Supplementary Fig. 6a). Next, we analyzed the functional gelatinase activity which would better reflect the overall proteolytic activity in the brain. Gelatinase activity in the ATT (*P* = 0.0026), ATT+SB-3CT (*P* = 0.0005) and ATT+Doxycycline groups (*P* = 0.0009) are significantly suppressed (Fig. 6d), and gelatinase activity is strongly associated with MMP-9 concentration (r = 0.902, *P* < 0.0001) (Fig. 6e). As expected, the decreased gelatinase activity is strongly associated with longer survival (r = -0.650, *P* < 0.0001) (Fig. 6f).

**Fig. 6.**
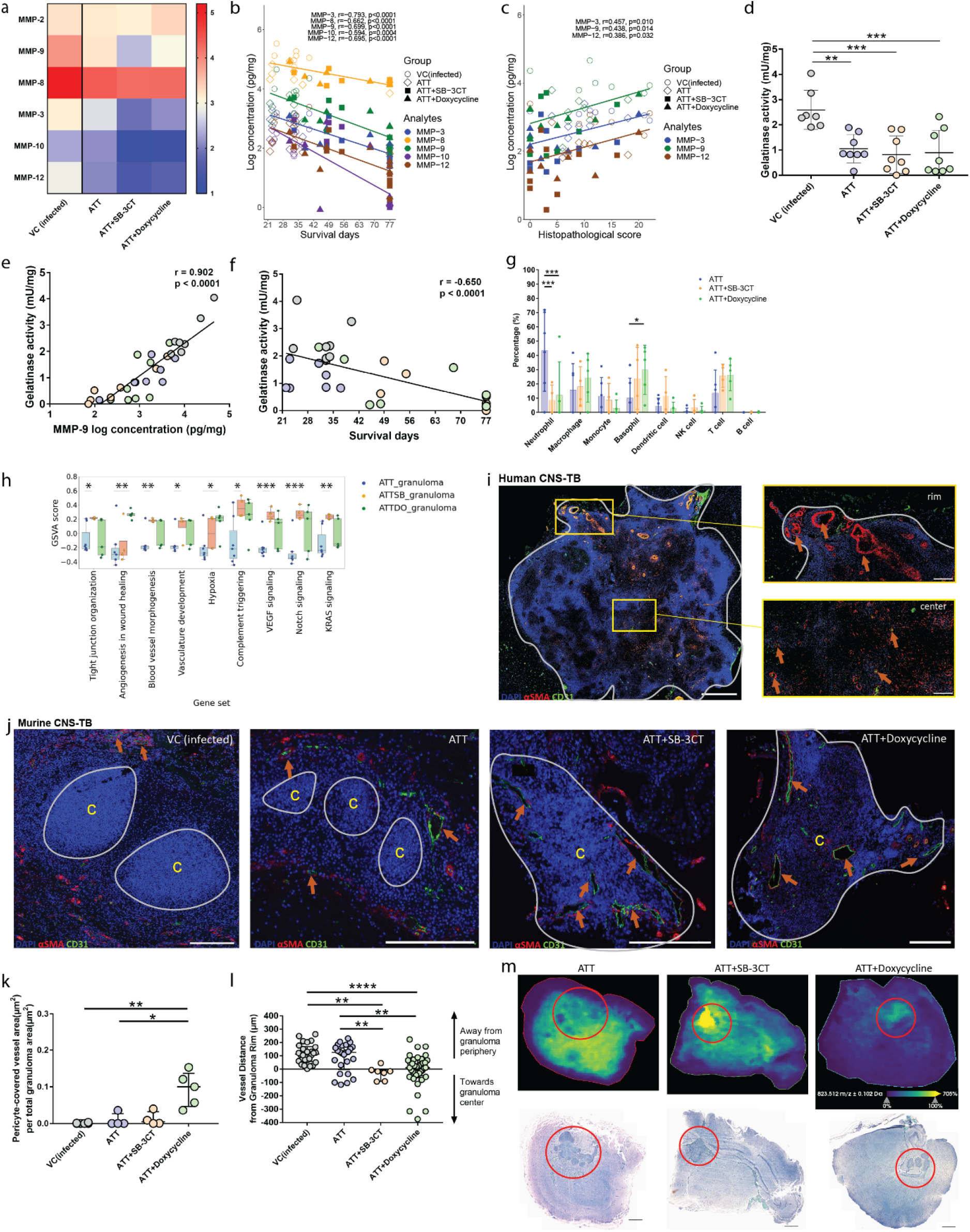
Adjunctive doxycycline with ATT suppresses functional gelatinase activity and normalizes blood vasculature in murine CNS-TB granulomas. **a**, Heatmap showing median expression of brain MMP-2, -9, -8, -3, -10, and -12 downregulated in the MMP inhibitor groups of murine CNS-TB. MMP concentrations are normalized to total protein concentration. **b**, Suppressed murine brain homogenate MMP-3, -8, -9, 10, and -12 are strongly associated with longer survival, analyzed by Spearman correlation coefficient. **c**, Up-regulated MMP-3, -9, and -12 are significantly associated with worse histopathological score, analyzed by Spearman correlation coefficient. **d**, Functional gelatinase activity is suppressed in the brain homogenates of murine CNS-TB which received ATT, ATT+SB-3CT or ATT+Doxycycline. Bars represent mean ± SD, analysis was conducted using one-way ANOVA with Dunn’s multiple comparisons test. **e,** Brain homogenate MMP-9 of murine CNS-TB is strongly associated with functional gelatinase activity, analyzed by Spearman correlation coefficient. **f,** The functional gelatinase activity of murine brain homogenates are strongly associated with longer survival, analyzed by Pearson correlation coefficient. g*, In silico* deconvolution analysis using ImmuCC signature matrix was conducted to infer immune cells fractions in murine CNS-TB of different treatment groups. Bars represent mean ± SD, and analysis was conducted using two-way ANOVA with Dunn’s multiple comparisons test. **h**, Enrichment of gene sets using the transcriptome data from murine CNS-TB treated with different regimes were analyzed by GSVA. Box represents 25th and 75th percentile, line represents median and whiskers denote the extremes. Analysis using unpaired t tests. * *P* < 0.05; ** *P* < 0.01, *** *P* < 0.001. **i,** Blood vessels were double-stained for endothelial cells with CD31and α-SMA for pericytes in human CNS-TB brain. Inset shows the granuloma rim and center. Vessels with wide lumen are distributed at the rim and mostly poorly-formed microvessels are found at the center. Gray outline indicates granuloma rim. Orange arrow points to blood vessel. Scale bar =200 µm. **j,** Compressed vessels and poorly formed microvessels are found in the granuloma periphery of *M. tb*-infected mice treated with VC or ATT alone. Conversely, vessels with open lumen are present in the granuloma center of *M. tb*-infected mice received adjunctive SB-3CT or doxycycline. Yellow “c” indicates granuloma center. Scale bar =200 µm. **k,** CNS-TB mice treated with ATT+Doxycycline have significantly increased total area of blood vessels with pericyte coverage compared to mice that received vehicle control or ATT alone. In each group, four to six tissue sections with presence of granulomas are assessed in a blinded fashion. Bars represent median ± IQR, analysis was conducted using Kruskal-Wallis test with Dunn’s multiple comparisons test. **l,** VC and ATT alone groups have more vessels located around the periphery, whereas the MMP inhibitor groups have more vessels distributed towards the center of the granuloma. Each point represents the distance of blood vessel from granuloma rim. The distance towards the granuloma center are calculated as a negative value. In each group, four to six tissue sections with seven to thirty-eight blood vessels within the maximum range of 300 µm away from granuloma rim was quantified. Bars represent median ± IQR, analysis was conducted using Kruskal-Wallis test with Dunn’s multiple comparisons test. **m,** Top panel shows representative MALDI-MSI images of murine CNS-TB brain tissues. Rifampicin is evenly distributed in the ATT alone group, in contrast to ATT+SB-3CT and ATT+Doxycycline groups which show higher concentrations within the granuloma. Bottom panel shows respective ZN stains. Red circled area indicates granulomas region with AFB stained. Scale bar =200 µm.

We performed LCM RNA-Seq to profile the transcriptomic changes in the brain tissues of murine CNS-TB treated with ATT, ATT+SB-3CT and ATT+Doxycycline. Low number of DEGs (*FDR* < 0.05) are identified (Supplementary Fig. 6b). The gene expression in murine CNS-TB that received different treatment regimes are heterogeneous (Supplementary Fig. 6c, Supplementary File 6). Based on the immune cells fractions inference from CIBERSORTx, the neutrophil fraction is significantly suppressed in the two groups of murine CNS-TB treated with adjunctive MMP inhibitors (ATT+SB-3CT, *P* = 0.0003; ATT+Doxycycline, *P* = 0.0005), while basophils are upregulated in murine CNS-TB on ATT with doxycycline (*P* = 0.0358) (Fig. 6g). Gene Set Variation Analysis (GSVA) analysis indicates that gene sets associated with tight junction organization (*P* = 0.0138), blood vessel morphogenesis (*P* = 0.0032), vascular development (*P* = 0.0436), complement (*P* = 0.022), VEGF (*P* = 0.0007), Notch (*P* = 0.0008) and KRAS (*P* = 0.0067) signaling are increased in murine CNS-TB on ATT with SB-3CT. Compared to ATT alone, ATT+Doxycycline shows increased expression in gene sets related to angiogenesis in wound healing (*P* = 0.0042) and hypoxia (*P* = 0.0139) (Fig. 6h, Supplementary File 7).

As we found MMP inhibition reduced AFBs in granulomas, reduced tissue destruction and enhanced gene sets associated with blood vasculature development, we sought to evaluate if the vasculature and drug distribution are also improved. In human CNS-TB brain tissue, well-formed vessels (CD31^+^αSMA^+^) with wide lumen area are found at the granuloma rim, whereas poorly-formed microvessels are located at the granuloma center (Fig. 6i). Similarly, murine CNS-TB in the vehicle control or ATT alone groups demonstrate blood vessels with compressed lumen along the granuloma periphery (Fig. 6j). In contrast, murine CNS-TB in the SB-3CT or doxycycline groups show blood vessels with pericytes (CD31^+^αSMA^+^) in both granuloma center and rim, with larger luminal area in ATT+Doxycycline group (Fig. 6j). Quantitatively, the ATT+Doxycycline group has significantly more well-formed pericyte-covered blood vessels compared to vehicle control (*P* = 0.0051) and ATT alone group (*P* = 0.0153) (Fig. 6k). Compared to ATT group, the well-formed pericyte-covered vessels are located significantly nearer to the granuloma center in ATT+SB-3CT (*P* = 0.0094) and ATT+Doxycycline (*P* = 0.0033) groups (Fig. 6l).

We also measured the drug distribution and concentration in the murine CNS-TB brains via MALDI-MSI and LC-MS/MS respectively. Comparable concentrations of ATT drugs Rifampicin, Isoniazid, Ethambutol and Pyrazinamide are detected in the brain homogenates for all the treated murine CNS-TB, indicating successful BBB penetration of ATT drugs in all the TB treatment regimes (Supplementary Fig. 6d). Doxycycline is also detected in the brain homogenates of mice received ATT+Doxycycline (Supplementary Fig. 6e, Supplementary Fig. 6f). Although we are unable to detect SB-3CT in the brain homogenates of *M. tb*-infected mice received ATT +SB-3CT, previous studies have proven the ability of SB-3CT to cross the BBB in mice^35, 36^. We further determined the distribution of rifampicin in the brain tissues using MALDI-MSI. Despite similar drug concentrations found in all treatment groups of murine CNS-TB (Supplementary Fig. 6d), rifampicin is homogenously distributed throughout the brain tissue in ATT-only group, indicating there was global capillary leak throughout the brain tissue, while with adjunctive MMP inhibition we observe more localized drug delivery (Fig. 6m). This may also result in more efficient *M. tb* killing thus explaining the lower AFB granulomas in the MMP inhibitor groups. Collectively, MMP inhibition in murine CNS-TB show suppressed MMPs which reduce tissue destruction and pathological lesions, thus decreasing both inflammatory neutrophil infiltration and NETs release. The less inflamed and non-necrotic tissues found in MMP inhibitor groups, with improved vasculature found in doxycycline-treated mice, enhances drug penetration into the granulomas to improve survival.

## Discussion

The current standard-of-care for human CNS-TB is ATT with steroids but this does not improve neurological outcomes for which new therapeutic strategies are urgently needed^37^. Here, we describe the crucial role of MMPs and NETs in human and murine CNS-TB, demonstrating their pathological role in driving CNS-TB immunopathology. Crucially, we found significant association of MMPs and NETs with neurological abnormalities and poor clinical outcomes. Spatial transcriptomic profiles of human and murine CNS-TB indicate a highly-inflamed and neutrophil-rich microenvironment, together with suppressed neuronal functions. In our pre-clinical drug trial, we demonstrate novel findings of ATT with doxycycline to improve murine CNS-TB survival and reduce immunopathology. We also uncover the mechanism of action of doxycycline and another MMP-inhibitor SB3CT, where blood vessel structure to the granulomatous regions of the brain is improved. Most importantly, our results are highly translatable for clinical use, as doxycycline is widely available and cheap which are key in TB endemic regions where resources are limited.

In the human cohort, compared to non-TBM patients, we found TBM patients to have significantly up-regulated CSF gelatinase MMP-2 and -9 and down-regulated TIMP-2 and -4, driving an ECM-degrading phenotype. Several human studies have previously demonstrated an upregulation of MMP-9 in TBM patients^2, 11–15^, which reflected the ability of *M. tb* to induce the secretion of human host enzymes to breakdown the basement membrane of the BBB^38^. The finding of increased CSF MMP-2 expression was similar in another pediatric TBM cohort^39^, but in contrast to previous findings in adult TBM^2^. We also found significantly higher expressions of MMP-1, -3, -7 and -10 in TBM patients who presented with neuroradiological abnormality and poor clinical outcomes, demonstrating that multiple MMPs contribute to the pathogenesis in CNS-TB. MMPs were reported to have pathological role in neurological illnesses such as stroke, multiple sclerosis and vascular cognitive impairment^40^. Furthermore, we describe novel findings of significantly increased CSF NETs in TBM patients which were associated with poorer neurological outcomes and death. Other non-TB studies had reported that NETs exacerbate neurological deficits in traumatic brain injury^41^ and poorer clinical outcomes in stroke^42^. In keeping with our findings in human CNS-TB, our pre-clinical CNS-TB murine model shows high expression of NETs and MMP-9 that colocalized at necrotic regions, implicating the critical role of NETs and MMP-9 in brain tissue destruction in this devastating infection.

Existing transcriptomic data on CNS-TB are limited to whole blood and CSF samples from TBM pediatric patients and brain tissues from *M. bovis*-infected mice^43, 44^. In this study, we present novel spatial transcriptomic analysis of the granulomas from human and murine CNS-TB using the *Nos2*^-^/^-^ strain. Our main findings reveal a significant enrichment of pathways associated with inflammation, immune responses, ECM organization and angiogenesis, with elevated neutrophil-related genes and TFs. Gene signatures from human TB lung tissues and whole blood have similarly shown that inflammation in TB are associated with interferon signaling pathways^45, 46^. We now extend findings of dominant IFN-ɣ and Type I IFN signaling which are neutrophil-driven in TB lungs^47^, to human and murine CNS-TB granulomas. Neutrophil qualitative states may be crucial determinants of disease^48^, thus we suggest to further characterize which neutrophil subsets are associated with IFN transcriptional program in CNS-TB using single-cell RNA sequencing. We also demonstrate that ECM remodeling of collagen and ECM degradation which is pathognomonic to CNS-TB are similar to the findings from TB lung and lymph node transcriptomic analyses^46, 49^. Down-regulated neuronal pathways in human and murine CNS-TB brain granulomas indicate that *M. tb* infection may interfere CNS neuronal function. The transcriptomic findings uncover a potential therapeutic intervention using MMP inhibitors to modulate host tissue remodeling and inflammation in CNS-TB granulomas.

In our CNS-TB mouse model^10^, we demonstrate adjunctive MMP inhibition reduces immunopathology and improves survival. We show MMP dysregulation in the brain of C57BL6 *Nos2^-/-^* mice, but not in C3HeB/FeJ mice, supporting the use of this mouse strain for CNS-TB. SB-3CT is a selective gelatinase inhibitor that crosses the BBB barrier^35^. Although previously used in a CNS-TB study, mortality benefit was not demonstrated^50^. Doxycycline, an U.S. Food and Drug Administration (FDA)-approved broad-spectrum MMP inhibitor is an effective adjunctive antibiotic which suppressed MMPs and reduced pulmonary cavity volume in pulmonary TB patients from our own Phase 2 trial^51^. Treating murine CNS-TB with ATT together with adjunctive SB-3CT or doxycycline, suppressed NETs, down-regulated MMP-3, -8, -9, -10 and -12 concentrations in the brain, with a corresponding reduction in CNS immunopathology, and resulted in significantly improved survival. Adjunctive MMP inhibition in our study also altered the immune cell fractions in murine CNS-TB, with significantly decreased neutrophils, increased basophils and a trend of increased T cells. Of note, previous studies have highlighted the importance of T cells in mediating *M. tb* control^52, 53^. In line with MMP inhibition which demonstrated increased pericyte-covered blood vessel and improved TB drug delivery and retention on murine pulmonary TB^54^, we now extend MMP inhibition to murine CNS-TB, and showed increased well-formed pericyte-covered blood vessels indicating that doxycycline modulates angiogenesis and normalizes vasculature in CNS-TB granulomas. Suppressing ECM remodeling may be the key mechanism by which MMP inhibition improves blood vasculature. MMP inhibitors markedly reduced collagen lysis, associated with stabilized endothelial networks with accumulation of collagen fibrils surrounding the microvessels^55^. As MMPs also detach pericytes from vessels during angiogenesis^56^, inhibiting MMPs increased pericyte coverage and attenuated vessel leakage^54, 57^. The modulation of blood vessels may facilitate drug penetration and retention in the granulomas, while concurrently suppressing inflammatory responses to hasten resolution of infection.

Our study has limitations. Given the rarity of CNS-TB, a proportion of our TBM patients had microbiologically-confirmed diagnosis, the remaining being possible and probable TBM^58^, but all patients received anti-TB treatment. With limited volume of CSF, we did not perform gelatin zymography to assess the endogenous activity of MMP-2 and -9. The human CNS-TB histological samples were archived up to six years, which may explain the lower gene counts compared to murine histological samples that were processed within a year. The transcriptomic data from murine CNS-TB mice of different treatments did not show clear clusters in PCA, which may be due to different harvesting point. The mice were not treated with steroids which is standard of care for human CNS-TB; we avoided steroids to determine the true effect of MMP inhibition. We were also constrained by the biosafety requirements to process *M. tb*-infected tissue as FFPE, and were unable to detect other TB drugs after the process of dewaxing the tissues.

Altogether, we demonstrate the detrimental role of MMPs and NETs in driving CNS-TB immunopathology both in a human cohort and in a pre-clinical murine model. Critically, adjunctive CNS-TB therapy with MMP inhibitors improve survival and reduce immunopathology in murine CNS-TB. Our findings show MMP inhibition has excellent potential to reduce the great toll of morbidity and mortality caused by CNS-TB, and doxycycline is readily available and cost-effective, as host-directed therapy in CNS-TB to improve both neurological and survival outcomes.

## Online Methods

### Ethics approval and consent to participate

The Domain Specific Review Board from National Healthcare Group Singapore (Reference: 2015/00067) approved the study. Anonymized human brain samples were used. All animal procedures were approved by the Institutional Animal Care and Use Committee of National University of Singapore under protocols R15-1068 and R21-0633, in accordance with national guidelines for the care and use of laboratory animals for scientific purposes.

### Participants

This was a prospective observational cohort study conducted at Hospital Likas, Kota Kinabalu, Sabah, East Malaysia. The study was approved by the Medical Research Sub-Committee of the Malaysian Ministry of Health (Reference NMRR-11-623-9629), the Menzies School of Health Research, Australia (Reference 2011-1636), and the Domain Specific Review Board, Singapore (Reference 2015/00067). Enrolment occurred from June 2013 till December 2014. A standardized data sheet was used by research clinicians to collect demographic data, epidemiological information (animal exposure, family history of TB, travel history), clinical history, vital signs, biochemical and microbiological data, imaging reports, treatment, outcome, and final diagnosis. Childhood vaccinations for potential CNS pathogens in Sabah, Malaysia included diphtheria-pertussis-tetanus, measles-mumps-rubella, polio, BCG, and *Haemophilus influenza type B* vaccine ^59^.

The inclusion of patients occurred in two stages. In the first stage, all pediatric patients with a clinical suspicion of meningitis or encephalitis were enrolled. At the second stage, the following case definitions for meningitis and encephalitis were used to exclude patients in the process. Meningitis was defined as having two or more of the following: i) fever or history of fever (≥38°C), ii) neck stiffness, iii) headache, or iv) cerebrospinal fluid (CSF) pleocytosis (≥ 5 WBC/µL). Encephalitis was defined as having encephalopathy (altered level of consciousness, lethargy, irritability, and change in behavior or personality) plus two or more of the following: i) fever or history of fever (≥38°C), ii) CSF pleocytosis (≥ 5 WBC/µL), iii) seizures or focal neurological findings, iv) abnormal neuroimaging consistent with encephalitis, or v) abnormal electroencephalogram findings compatible with encephalitis. This definition had been used previously and will allow us to compare our results to published studies^59^. A published consensus case definition was used to classify TBM into confirmed, probable or possible TBM based on clinical, CSF, microbiological and radiological criteria^58^. Clinically diagnosed and treated TBM includes probable (n = 9) and possible (n = 9) cases.

Exclusion criteria were (i) patients with non-infectious CNS disorders due to hypoxic, vascular, toxic and metabolic causes, (ii) patients with CNS disorders lasting less than 24 hours, and (iii) patients with malaria diagnosed on microscopy. Patients with an immune-mediated post-infectious etiology such as acute disseminated encephalomyelitis (ADEM) were enrolled as well.

### Clinical neurological scoring, radiological abnormalities and outcomes assessment

The modified Glasgow Coma Scale (GCS) was used to assess the conscious state of patients ≥ 3 years old, with scores ranging from 3 (indicating deep unconsciousness or coma) to 15 (fully awake). The scale is composed of 3 tests: eye response (score 1-4), verbal response (score 1-5) and motor response (score 1-6). In younger children (patients < 3 years old), the Blantyre Coma Scale (BCS) was used instead, which scores from 0-5 with 0 being equivalent to GCS 3 and 5 being equivalent to GCS 15. Similar to GCS, the BCS is also composed of 3 tests: eye response (score 0-1), verbal response (score 0-2) and motor response (score 0-2). CT brain of TBM patients were assessed for radiological abnormalities including leptomeningeal contrast enhancement and ventricular dilatation by a neuroradiologist blinded to patient outcomes. TBM patients with full recovery or good recovery with mild deficit are categorized under good clinical outcome, whereas TBM patients who died or survived with severe disability were categorized under poor clinical outcome.

### Mouse cannula implantation and infection

Six-to-eight-week-old male C57BL/6 *Nos2^−/−^*(Stock No. 002609) and C3HeB/FeJ mice (Stock No. 000658) (Jackson Laboratory) were infected with *M. tb* H37Rv at seven days after brain cannulation, via intra-cerebroventricular (i.c.vent.) route as previously described^10^. Briefly, a motorized stereotaxic instrument (Neurostar) was used to implant a 26-gauge guide cannula (RWD) into the third ventricle (coordinates from the bregma: − 1.6 mm posterior, 0 mm lateral, − 2.5 mm ventral) of mice at one week before infection. Mice were injected with 0.5 μL of sterile 0.9% NaCl or 2 × 10^8^ CFU/mL *M. tb* (administered at a dose of 10^5^ CFU to each animal) through the brain cannula (over 5 min) using the syringe pump (Harvard Apparatus). All mice were observed for mortality and weight change. Humane endpoints included ⩾ 20% weight loss, complete hind limb paralysis and repeated seizures. Infected mice were also monitored daily after infection for clinical signs indicative of CNS-TB, such as limb weakness, tremors, and twitches.

### Drug treatment for *M. tb-*infected mice

Anti-tuberculous (ATT) drugs Isoniazid (INH), Rifampicin (RIF), Pyrazinamide (PZA), and Ethambutol (EMB) were purchased from Sigma. All drugs were administered five days a week for a total duration of eight weeks via oral gavage in the following dosages as previously used in other murine TB studies; RIF(10 mg/kg), INH(25 mg/kg), PZA(150 mg/kg), and EMB(100 mg/kg)^60, 61^. Drug solutions were prepared such that the desired concentration would be delivered in a 0.2 mL total volume. The 25 mg/kg dose of INH produces a serum area under the concentration-time curve from 0 to 24 h (AUC_0-24_) value similar to those observed in humans with a slow acetylator phenotype^62^. RIF was given 45 min prior to administration of the other drugs to avoid any adverse pharmacokinetic interaction^63–65^.

SB-3CT (MedChemExpress) (50 mg/kg) or doxycycline (Sigma) (100 mg/kg) were administered into mice by intra-peritoneal (i.p.) injection for five days per week for a total duration of eight weeks. Based on allometric scaling calculation^66^, the human equivalent dose for doxycycline use here is 8.1mg/kg, which exceeds the maximum clinical dose for doxycycline at 100 mg twice daily. However, it has previously reported that C57BL/6 mice given oral doxycycline dose of 100mg/kg daily had similar mean plasma concentration of doxycycline as patients treated with 200 mg doxycycline daily^67^, thus we used a similar dose in this study. SB-3CT or doxycycline was diluted in the diluent comprises of 10% DMSO, 75% PEG-200 and 15% milli Q water daily prior to i.p. injection. Drug suspension was prepared such that the desired concentration would be delivered in a 50 µL total volume.

### Histopathological analysis

Histopathology was performed on the left hemisphere of the murine brain, which was fixed in 10% neutral buffered formalin, paraffin-embedded and sectioned to 4 μm thickness. Brain slices were stained with hematoxylin–eosin (Vector Laboratories, Burlingame, California) to characterize pathological lesions and Ziehl–Neelsen staining (Sigma-Aldrich, St. Louis, Missouri) to detect mycobacterium according to manufacturer’s instructions. Histopathology was evaluated in a blinded fashion by a histopathologist (R.R.) based on the presence of pathological changes including inflammation, perivascular cuffing, gliosis, granuloma (not well developed), pyogranuloma, and neuronal degeneration. Grading of severity was assigned on the following scale: 0: no abnormalities detected; 1—minimal; 2—mild; 3—moderate; 4— marked and 5—severe. The presence of liquefactive necrosis and presence of bacilli were also accessed.

### Organ harvesting and processing

Mouse brain was harvested at eight weeks after infection, or eight weeks post-treatment for mice in the drug study. Half of the brain was fixed in 10% neutral buffered formalin for histology, and the other half was homogenized for bacterial enumeration and characterization of immunological markers. The organ was homogenized by high-speed shaking in 2 mL microcentrifuge tubes filled with sterile PBS and 5/7 mm stainless steel beads using TissueLyser LT (Qiagen, Hilden). For bacterial enumeration, brain homogenates were cultured on selective Middlebrook 7H11 agar supplemented with 0.05 mg/mL carbenicillin (Sigma), 0.01 mg/mL amphotericin B (Sigma), 0.02 mg/mL trimethoprim (Sigma), 200 units/mL polymyxin B (Sigma), 0.5% glycerol and 10% OADC.

### Sterilization of human CSF and murine brain homogenates

Following experiments involving *M. tb*, samples required removal of *M. tb* by double filtration before analysis. CSF and organ homogenates are transferred to 0.22 µm centrifugal filter units (Millipore) and centrifuged at 12,000 x g for 5 min. The filtrate is again centrifuged using a new filter unit so each sample will be sterile filtered twice.

### Total protein concentration quantification

Total protein concentration was quantified using DC assay (Bio-Rad) according to manufacturer’s instructions. Five microliters of standard or samples were loaded into 96-well plate. Next, 25 µL of Reagent A was added in each well, followed by additional 200 µL of reagent B added into each well. The plate was placed on shaker and mixed for 5 sec. After 15 min incubation at room temperature, the absorbance was read at 750 nm using a microplate reader.

### Luminex bead array for human MMPs and TIMPs

Human CSF MMP-1, -2, -3, -7, -9, -10 and EMMPRIN concentrations were analyzed by Fluorokine multianalyte profiling kit (R&D Systems). Human CSF TIMP-2 and -4 concentrations were analyzed by human TIMP multiplex kit (R&D Systems). All analytes were analyzed using the Bio-Plex 200 platform (Bio-Rad), according to the manufacturer’s protocol. The minimum detection limit for the nine analytes were 1.1, 12.6, 7.3, 6.6, 13.7, 3.2, 5.6, 14.7, 1.29 pg/mL, respectively. CSF samples were diluted with calibrator diluent with 1 in 5 dilution for the array of MMPs, and 1 in 100 dilution for the array of TIMP-2. Neat CSF was used to analyze TIMP-4 concentration. Twenty-five microliters of standard or samples were loaded into 96-well plate, and incubated with 25 µL of microparticle cocktail (diluted 100-fold with Microparticle Diluent) at room temperature for 2 h shaking at 800 rpm. The microplate was washed three times, 25 µL of diluted Biotin Antibody cocktail was added and the plate was incubated for 1 h on the shaker. After three washes, 25 µL of streptavidin-PE was added, incubated for 30 min and further washed three times. The microparticles were resuspended with 100 µL of wash buffer, incubated for 2 min on the shaker and read using the Bio-Plex analyzer. CSF concentrations of analytes were normalized to total CSF leukocyte count. For patients who have 0 leukocytes in the CSF, the MMP concentration was normalized to 1 leukocyte.

### Luminex bead array for mouse MMPs, adhesion molecules, cytokines and chemokines

Analysis of mouse biomarker concentrations were performed using the Mouse Premixed Multi-Analyte Kit (R&D Systems) according to manufacturer’s instructions. The analytes with respective minimum detection limit are MMP-2 (48.8 pg/mL), MMP-3 (0.332 pg/mL), MMP-8 (2109 pg/mL), MMP-9 (47.1 pg/mL), MMP-12 (0.424 pg/mL), TIMP-1 (12.4 pg/mL), TIMP-4 (14.4 pg/mL), TNF-α (1.47 pg/mL), IFN-ɣ (1.85 pg/mL), IL-1α (8.17 pg/mL), IL-1β (41.8 pg/mL), IL-6 (2.3 pg/mL), IL-17A (7.08 pg/mL), IL-27 (1.84 pg/mL), CXCL1 (32.9 pg/mL), CXCL2 (1.97 pg/mL), CXCL10 (6.85 pg/mL), CXCL13 (19.3 pg/mL), CCL2 (134 pg/mL), CCL3 (0.452 pg/mL), CCL4 (77.4 pg/mL), CCL5 (19.1 pg/mL), CCL7 (1.69 pg/mL), CCL11 (1.46 pg/mL), CCL12 (0.613 pg/mL), CCL19 (2.28 pg/mL), and CCL22 (0.965 pg/mL). Neat concentrations of mouse brain homogenates were used. Twenty-five microliters of standard or samples were loaded into 96-well plate, and incubated with 25 µL of microparticle cocktail (diluted 10-fold with Assay Diluent RD1W) at room temperature for 2 h shaking at 800 rpm. The microplate was washed three times, 25 µL of diluted Biotin Antibody cocktail was added and the plate was incubated for 1 h on the shaker. After three washes, 25 µL of diluted streptavidin-PE was added, incubated for 30 min and further washed three times. The microparticles were resuspended with 100 µL of wash buffer, incubated for 2 min on the shaker and read using the Bio-Plex 200 analyzer.

### Gelatinase inhibition assay using MMP-2 and MMP-9 antibodies

Gelatin degradation by neat concentration of CSF was assessed. Samples were activated with 2 mM of aminophenylmercuric acetate (APMA) for 2 h at 37°C. Next, 80 ng of inhibitor MMP-2 antibody (Thermo Fisher MA513590) or MMP-9 antibody (Sigma-Aldrich IM-09L) was added per well and incubated for 1 h at 37°C. Subsequently, 1 µg of DQ gelatin was added per well and incubated for 6 hours in the dark at room temperature. Gelatin degradation activity was measured at absorption 485 nm and emission 528 nm with a fluorescence plate reader (BioTek Instruments). Results were normalized by the total protein concentration.

### DQ gelatin degradation assay

Gelatin degradation by neat concentration of murine brain homogenates was assessed. Samples were activated with 2 mM of aminophenylmercuric acetate (APMA) for 2 h at 37°C, before mixing with 1 µg of DQ gelatin. After 16 hours incubation in the dark at room temperature, gelatin degradation activity was measured at absorption 485 nm and emission 528 nm with a fluorescence plate reader (BioTek Instruments). Results were normalized by the total protein concentration.

### NETs quantification

NETs concentrations were quantified by measuring extracellular DNA concentration, a commonly used method to evaluate NETs levels in both *in vitro* and clinical studies^9, 68, 69^. To take into account the number of neutrophils entering the CNS, extracellular DNA concentrations were corrected for CSF neutrophil count. NETs were quantified using QuantiT PicoGreen (Invitrogen) according to manufacturer’s instructions. In brief, a standard was made using serial 1 in 10 dilutions of Lambda DNA commencing from 20 µg/mL to 2 ng/mL diluted in 1 X TE buffer. Fifty microliters of samples or standards were loaded onto 96-well black plates, and 50 µL PicoGreen at a 1 in 200 dilution was added in each well. The plate was then incubated in the dark at room temperature for 5 min, and then read using the Synergy H1 microplate reader (BioTek).

### Enzyme-linked immunosorbent assay (ELISA) for Citrullinated Histone 3 (CitH3)

CitH3 concentration was assayed using Citrullinated Histone 3 (Clone 11D3) ELISA kit (Cayman Chemical), according to the manufacturer’s instructions. CSF or brain homogenates were diluted 1 in 2 with Assay Buffer. Serial 1 in 2 dilutions of standards were made commencing from 10 to 0.15 ng/mL. One hundred microlitres of standards or samples were added per well and incubated for 2 h at room temperature on a horizontal orbital microplate shaker. The plate was washed five times with Wash Buffer before adding 100 µL of 1X HRP Conjugate per well, and the plate incubated for another hour at room temperature. The plate was washed five times, prior to the addition of 100 µL of TMB Substrate Solution and incubated for 30 min in the dark at room temperature. One hundred microliters of Stop Solution was added to each well, and the optical density was read using a microplate reader set to 450 nm. The lower limit of detection is 0.1 ng/mL.

### *Mycobacterium tuberculosis* culture for infection

*M. tb* strain H37Rv was kindly provided by Professor Nicholas Paton (National University of Singapore, Singapore). *M. tb* was cultured in Middlebrook 7H9 medium (BD Difco) supplemented with 10% (v/v) Middlebrook albumin-dextrose-catalase (ADC) (BD Difco), 0.2% (v/v) glycerol (Sigma) and 0.05% (v/v) Tween-80 (Sigma) or on complete Middlebrook 7H11 agar (BD Difco) supplemented with 0.5% (v/v) glycerol and 10% (v/v) oleic acid-albumin-dextrose-catalase (OADC) (BD Difco). *M. tb* cultures were sustained in square PETG bottles (Nalgene) at 37 ᴼC with agitation at 80 rpm or plated on complete or selective 7H11 agar in petri dishes (Thermo Fisher Scientific). For infection experiments, *M. tb* was cultured to mid-logarithmic phase at an optical density of 0.6 to 0.8. Prior to infection, the *M. tb* was centrifuged at 3200g for 10 min and resuspended in 1 mL sterile 0.9% NaCl. The *M. tb* inoculum was then plated to determine the dose of infection.

### ELISA for mouse TIMPs

Mouse TIMP-2 concentration was measured using the Duoset ELISA Development System (R&D Systems) according to the manufacturer’s instructions. The lower limit for detection of TIMP-2 is 7.8 pg/mL. ELISA plates were coated with 100 µL of Capture Antibody at 1 µg/mL overnight at room temperature, then washed three times with Wash Buffer (0.05% Tween-20 in PBS). Free binding sites were blocked for 1 h with 300 µL of Reagent Diluent (1% BSA in PBS) at room temperature, then washed three times. Fifty µL of standards or brain homogenates (diluted 1 in 100 to 200 with water) were added to the appropriate wells and incubated for 2 h at room temperature. After three washes, 50 µL of Detection Antibody (50 ng/mL) was added for 2 h at room temperature. Three further washes were performed, then 50 µL of Streptavidin-HRP (1 in 40 dilution) was added into each well and incubated for 20 min in the dark at room temperature, before washing three times again. Fifty µL Substrate Solution (1:1 mixture of H_2_O_2_ and Tetramethylbenzidine) was then added per well and the plate incubated for 20 min in the dark at room temperature. The reaction was then stopped with 25 µL per well of Stop Solution (2 N H_2_SO_4_) and the plate read at 450 nm and 570 nm using the Synergy H1 microplate reader (BioTek, US). The 570 nm reading was subtracted from the 450nm reading to account for imperfections in the plate.

Mouse TIMP-3 concentration was measured using ELISA kit (LSBio), according to the manufacturer’s instructions. The lower limit for detection of TIMP-3 is 0.156 ng/mL. Brain homogenates were diluted 1 in 50 with water. One hundred microliters of standard or sample was added per well and incubated for 2 h at 37 ᴼC. After 2 h incubation, mixture of each well was aspirated without washing. One hundred microliters of 1X Biotinylated Detection Antibody was added into each well, incubated for 1 h at 37 ᴼC, then washed three times. Each wash step was allowed to sit for 2 min before completely aspirating. One hundred microliter of 1X HRP-streptavidin was added per well and incubated for a further hour at 37 ᴼC. The plate was washed five times, prior to the addition of 90 µL of TMB substrate and incubated for 15 to 30 mins in the dark at 37 ᴼC. Fifty microliters of stop solution was added to each well, and the optical density was read using a microplate reader set to 450nm and 570nm. The 570nm reading was subtracted from the 450nm reading to account for imperfections in the plate.

### Immunohistochemistry

Tissue sections were deparaffinized and rehydrated with 5 min each in sequentially order of Clearene, Clearene, 100% Ethanol, 100% Ethanol, 95% Ethanol, 70% Ethanol, and distilled water. Subsequently, heat-induced epitope retrieval was performed by incubating the sections in 1x Citrate Buffer, pH 6.0 (Sigma), for 30 min at 95 ᴼC. After antigen retrieval, sections were left in the Citrate Buffer at room temperature to cool for 20 mins, before washing with PBS three times with 5 min each. Sections were surrounded with PAP pen (Vector laboratories) and treated with blocking buffer for 1 h at room temperature to prevent non-specific binding. Blocking buffer used was 10% goat serum in PBS. After blocking, primary antibodies were added to the sections. The antibodies with respective incubation condition are anti-histone H3 (citrulline R2 + R8 + R17) antibody (Abcam Ab5103, 1:50 dilution, 37C, 2 h incubation), anti-MMP9 antibody (Abcam Ab38898, 1:250, RT, 2 h incubation). Blocking buffer alone was used as a control. Upon primary antibody incubation, sections were washed three times with PBS for 5 min prior to the addition of secondary antibodies ImmPRESS goat anti-rabbit-HRP (Vector Laboratories MP7451) for 30 min incubation at RT. All primary and secondary antibodies were diluted in 10% goat serum in PBS.

For chromogenic staining, the DAB Substrate Kit (Abcam) was used according to the manufacturer’s instructions. Sections were covered with DAB Substrate for 1 to 10 min until the desired color is achieved, then washed under running tap water for 2 min before counterstaining with hematoxylin for 5 min. Sections were washed with tap water again for 2 min followed by dehydration with 3 min each in sequentially order of 50% Ethanol, 70% Ethanol, 90% Ethanol, 100% Ethanol, 100% Ethanol, Clearene, and Clearene. Histological sections were covered with Organo/Limonene Mount and coverslip, air-dried overnight and then stored at room temperature. Negative control for all IHC stains showed no immunoreactivity.

For double immunofluorescence staining involving the use of two primary antibodies raised in the same host, Opal 3-Plex Anti-Rb Manual Detection Kit (Akoya Biosciences) were used according to the manufacturer’s instructions. After dewaxing and rehydration, antigen retrieval of sections was performed using microwave treatment, 1 min at 100% power followed by 15 min at 20% power. To block endogenous HRP activity, sections were incubated in 1% H2O2 in TBS for 1 hour and washed, followed by blocking in 10% goat serum in TBS. All washing step includes washing in 1X TBS-Tween 20 (0.05%) for three times in 5 min each. Anti-CD31 antibody (Abcam Ab182981) was added to the sections at 1:500 dilution, for 16 hours incubation at 4°C. After washing, secondary anti-rabbit HRP from the kit was added for 15 min incubation at RT. Sections were washed, then incubated with Opal 520 dye for 15 mins at room temperature and further washed. Prior to the addition of the second primary antibody, the first primary antibody was eluted by repeating the microwave treatment. After washing, anti-αSMA antibody (Abcam Ab124964) was added to the sections at 1:500 dilution, for 16 hours incubation at 4°C. After washing, secondary anti-rabbit HRP from the kit was added for 15 min incubation at RT. Next, the sections were incubated with Opal 570 dye for 15 mins at room temperature. After washing, nuclei were counterstained with DAPI solution for 5 mins before the final washing step. The sections were mounted in ProLong Gold Anti-fade aqueous mounting medium. Negative control for all IHC stains showed no immunoreactivity.

### Laser capture microdissection RNA sequencing

Archival FFPE (formalin-fixed paraffin-embedded) tissue from patients with clinical biopsies for CNS-TB and patients with demyelination as controls were obtained from the clinical pathology unit with ethics approval. For murine samples, TB-infected mouse brains and controls were fixated in formalin and embedded in paraffin blocks.

Paraffin sections of 7µm thickness were mounted on polyethylene naphthalate (PEN) membrane slides for laser-capture microdissection, with an adjacent H&E section used to confirm the region of interest. PEN membrane slides were deparaffinized with a series of xylene and ethanol wash, then stained with hematoxylin (Dako #S3309) and counterstained with bluing reagent (Thermofisher Catalog #6769002). Laser-capture microdissection was performed on the Leica LMD6 system, targeting regions of interest, including: granulomas, regions of inflammation, and areas of healthy tissue as controls. Biological duplicates were collected in separate tubes. RNA-Seq libraries were prepared using Smart-3SEQ^70^. Libraries were evaluated for their fragment size using the Agilent TapeStation4200 and quantified by qPCR on the Applied Biosystem StepOne Plus real-time PCR machine. Libraries were pooled and sequenced on an Illumina NextSeq500.

### Bioinformatics analysis

Quality control checks of raw sequencing data were executed using FastQC software (version 11.9). Alignment was performed using Kallisto (version 0.46)^71^, with sequence based bias correction. The alignment was performed against the human transcriptome (*Homo sapiens* GRCh38), mouse transcriptome (*Mus musculus* GRCm39), and *M. tb* strain H37Rv transcriptomes, which served as index transcriptomes. Data was annotated using ensembldb package (version 2.24.0). The transcript abundance was summarized to gene level using Sleuth (version 0.30)^72^. Raw counts from RNA sequencing were processed using edgeR, Bioconductor version 3.17^73^, with 3 variance estimated and size factor normalized using Trimmed Mean of M-values (TMM).

Differential analysis was performed using edgeR (version 3.17) and filterByExpr function was used to filter genes that have low number of reads. Samples with median that is beyond twice the median absolute deviation (MAD) from the median of medians, and any observations that are more than 2 IQR below Q1 or more than 2 IQR above Q3 are considered outliers were excluded from downstream analysis. Design matrix coded by model.matrix(∼0+variable) was used to identify the DEGs comparing two groups of interest. Genes with a false discovery rate (*FDR*)-corrected p-value < 0.05 are considered differentially expressed, following on from a likelihood ratio test using a negative binomial generalized linear model fit. Pathway enrichment analysis of significant up- and down-regulated DEGs was performed using web-based ShinyGO v0.77^74^. Limma with its voomWithQualityWeights function (version 3.56.2) was performed and the counts were used for principle component analysis. Gene signature enrichment was performed using Gene Set Variation Analysis (GSVA)^75^ using the gene sets from Molecular Signatures Database (MSigDB) (v2023.1)^76^. Key transcription regulators and their target genes are analyzed using web-based TTRUST v2^77^.

*In silico* cell fraction deconvolution was performed with web-based CIBERSORTx to estimate the abundance of immune cell types. LM22 signature matrix which distinguishes 22 purified immune cell types is used for human dataset^30^, and ImmuCC signature matrix is used to infer the relative proportions of 10 major immune cells for mouse dataset^78^.

### Granuloma and blood vessel analysis

The brain tissues harvested from *M. tb* H37Rv-infected C57BL/6 *Nos2^-/-^* mice treated with ATT alone, or ATT with adjunctive MMP inhibitors, or vehicle control (n = 7 to 8 per group) were processed as FFPE sections. The first, tenth, twenty-fifth and fiftieth sections away from *M. tb* infection site were stained with hematoxylin-eosin. The number of granulomas present in each section was quantified manually in a blinded fashion by H.T.C. and T.H.H. Ziehl– Neelsen staining was performed on adjacent tissue sections and evaluated for the presence of AFB. Full slide HE and ZN stains were scanned using TissueFAXS Slide Scanner (TissueGnostics). The section presented the most severe histopathology was selected as the representative section for each group, and the adjacent sections were used for vasculature analysis and drug distribution analysis.

Blood vessels of both human and murine CNS-TB tissues were double-stained with antibodies detecting CD31 for endothelial cells and α-SMA for pericytes, which maintain the integrity of vessels and the BBB^79^. IHC stains were scanned using Vectra 2 (PerkinElmer). Blood vessels were analyzed using customized semiautomatic software applications (Visiopharm). Deep learning application was trained for automatic selection, followed by manual verification and modification to generate outputs. The granulomas areas were outlined manually as region of interest (ROI). Within the ROI, area with CD31 stain was quantified as vessel area, while area with CD31 and αSMA double stains were quantified as vessel area with pericyte coverage. The distance of blood vessels from granuloma rim within the maximum range of 300 µm away from granuloma rim was quantified using ImageJ software.

### Liquid chromatography-tandem mass spectrometry

Twenty-five microliters of mouse brain homogenates and standard calibrators were transferred into a deep well 96-well plate and treated with 180 μL of ice-cold Acetonitrile containing 0.1% Formic acid. The plate was mixed on a shaker at 1000 rpm/min for 10 min, then centrifuged at 2270 x g for 50 min at 4 °C. One hundred microliters of the supernatant was carefully transferred to a 96-microwell plate and loaded into the auto-sampler (7 °C) for analysis by LC-MS/MS. Ion counts or areas under the MRM chromatogram were utilized for the construction of a standard curve for partial quantification.

Multiple reaction monitoring (MRM) methods, analyses, and data acquisition using Agilent’s 1290 Infinity II LC/ 6495 tandem triple quadrupole mass spectrometer equipped with electrospray ionization (ESI) source in MRM mode and HILIC chromatographic conditions were as follows: Liquid chromatographic (LC) separation of the biomarkers was carried out on an Agilent 1290 Infinity II LC system (Agilent Technologies) with PEEK coated SeQuant®ZIC®-cHILIC 3mm,100Å 100 x 2.1 mm HPLC column (Merck Pte Ltd, Singapore) maintained at 40 °C. The organic solvent was Methanol containing 2 mM Ammonium Formate and 2% Formic acid (Solvent A) and the aqueous solvent used was water containing 2 mM Ammonium Formate and 2% Formic acid (Solvent B). A linear LC gradient on Binary Pump (Agilent Model G7120A) was set up with a percentage of Solvent B as follows: 60% between 0 and 0.05 min, 10% between 1.00 and 2.00 min, and back to 60% between 2.01 and 3.5 min, with a flow rate of 0.4 mL/min. The column was further equilibrated for a further 7 min with 60% Solvent B. The sample injection volume was 10 μL. For mass detection, the LC eluent was connected online to an Agilent 6495 Triple Quadrupole MS system (Instrument model G6495A, Agilent Technologies) operated with the electrospray source in positive ionization mode. The electrospray ionization source conditions were as follows: capillary voltage of 4.0 kV, nozzle voltage of 500 V, iFunnel parameter high/low-pressure RF of 200 V, nebulizer pressure of 45 psi, the gas temperature of 290 °C, sheath gas temperature of 290 °C, and sheath gas flow of 20 L/min. For quantitative analyses, the analytes were detected by means of characteristic product ions formed from protonated molecules by collision-induced dissociation (CID) using the MRM mode after fragmentation with 99.9995% pure nitrogen. The MRM transitions were as shown in Supplementary Table 4. Dwell time was set to 100 ms and all transitions were in positive mode. Acquisitions and quantitative analysis were performed using the Agilent MassHunter Workstation Acquisition (10.0.127) and Agilent MassHunter Workstation Quantitative Analysis (10.0) software.

Calibration curves were constructed using standard stock solutions in water. For quantification, a calibration curve for each compound was constructed by serial dilution at decreasing concentrations. QCs were set at low, mid, or high concentrations. A linear curve was plotted, and the limits of quantitation (LOQ) were determined. The linearity coefficients of determination (r2) for all metabolites were > 0.9.

### Mass spectrometry imaging

The section presented severe histopathology with the presence of granulomas was selected as the representative section for each mouse group. FFPE tissue sections (5µm thickness) embedded on the conductive glass slide, Intellislide (Bruker Corporation) underwent dewaxing process. The slides were placed in an oven at 60°C for 60 mins to melt the parafilm. Then the slides were immersed in solution in the following order to remove the parafilm: 100% xylene twice for 3 mins each time, 100% EtOH twice, 95% EtOH once, 70% EtOH once, for 1 min each time, and lastly in water twice for 3 mins each time. The slides were placed in a vacuum desiccator for about 15 mins to dry.

After drying, external calibrants of poly-DL-alanine (pA) (Sigma-Aldrich) (0.15 mg/mL in H2O) and a mixed standard of INH, PZA, EMB, RIF, Doxycycline, and SB-3CT (2.5 to 25 µg/mL in H2O, stock SB-3CT in DMSO) were spotted (0.5 µL) at two locations on each slide and dried again in the vacuum desiccator. The slides were then scanned with the flatbed scanner Epson Perfection V39 (Nagano) at 3200 dpi. After scanning, the slides were sprayed with HCCA solution (Bruker Corporation) (10 mg/mL in 70%/30%/0.2% ACN/H2O/TFA) with the M3 TM sprayer (HTX Technologies LLC) with solvent flow of 0.12 mL/min, nozzle temperature of 75°C, four passes in horizontal spray pattern, 1200mm/min nozzle velocity with a track spacing of 3mm. After spraying, the slides were stored in 50 mL Falcon tubes with silica gel as desiccants until MSI experiment.

The tissue sections were scanned with the rapifleX TOF/TOF MALDI mass spectrometer (Bruker Daltonik). The mass spectrometer was operated in positive, reflector mode. External mass calibrations were performed using spots of pA and mixed standard spotted on the slide. Mass spectra were acquired in the mass range of 100 to 1000 Da, at a spatial resolution of 50 µm, with the Smartbeam 3D laser operated in Single beam mode at 40% laser fluence power. A blank area of matrix and a small mixed standard section (<200 positions) were also scanned.

Post-acquisition analysis was done using the SCiLS Lab 3D (v. 2016b) software (Bruker Daltonik). Root mean square (RMS) algorithm was used for normalization. The TB drug ions within the tissue sections data were annotated by matching ion peaks with the previously identified mixed standards peaks. Automatic hotspot removal, and weak denoising were used for ion image processing.

### Statistical analysis

The statistical analyses performed for continuous variables depended on the distribution (parametric or nonparametric) of the data. Where the data followed a normal distribution (based on D’Agostino-Pearson or Shapiro-Wilk normality test), the Student’s t-test for two groups or one-way ANOVA with Dunnett’s multiple comparison test against control group was used; Mann-Whitney or Kruskal-Wallis with Dunn’s multiple comparisons test was used for data with non-normal distribution. In correlation analysis, normally distributed data was analyzed with Pearson correlation coefficient, and non-normal distributed data was analyzed with Spearman correlation coefficient. The log-rank test was used for the analysis of Kaplan-Meier survival curves. Data are presented as median and interquartile range or mean and standard deviation as stated in the figure legend. A two-sided value of p < 0.05 was considered statistically significant. All analyses were performed using GraphPad Prism version 7.05 (GraphPad).

## Reporting Summary

Further information on research designs is available in the Nature Portfolio Reporting Summary linked to this article.

## Data availability

Demographic data of pediatric patients have been made available in the Supplementary. The raw sequence data generated in this study was deposited at Gene Expression Omnibus (GEO) repository under the accession number GSE244089.

## Code availability

The scripts used for the RNA-sequencing analysis are available through https://github.com/feikeanloh/CNSTB.

## Acknowledgements

The work was funded by NUHSRO/2016/066/NPCseedfunding/01 and NMRC/TA/0042/2015. C.W.M.O. is funded by Singapore National Medical Research Council (NMRC/TA/0042/2015, CSAINV17nov014, CSAINV21nov-0003; National University Health System Singapore (NUHS/RO/2017/092/SU/01,), and iHealthtech at the National University of Singapore. X.Y.P. was supported by a postgraduate scholarship from the Yong Loo Lin School of Medicine, National University of Singapore. F.K.L. and J.K.T. are funded by the NUSMed B2B Collaboration Grant by NUS Yong Loo Lin School of Medicine and NUHS Research Office. F.K.L. received NCID Catalyst Grant and NCID Short Term Fellowship administered by the National Centre for Infectious Diseases. J.K.T. is funded by the Singapore National Medical Research Council (NMRC/TA/20nov-0025 and NMRC/OFLCG/21Jun-0013). J.M.H. and F.K.L. were supported by NUSMed Post-Doctoral Fellowship (NUHSRO/2018/052/PDF/04). Q.H.M. and C.B. were supported by NUSMed Post-Doctoral Fellowship (NUHSRO/2017/073/PDF/03). S.C. received NUHS Junior Research Award and Chan Heng Leong Education and Research Fund. B.V. was supported by National Science Foundation HRD-1202008 and HRD-1826745.

We thank the operations team of the National University of Singapore BSL-3 core facility for the infrastructure and logistical support of the study. We express gratitude to National University of Singapore Comparative Medicine (CM) for animal training and support. We are thankful to the Microscopy and Multiplex Assays Core at the Cancer Science Institute and Dr Anand D Jeyasekharan for Visiopharm training and application. We also thank the Confocal Microscopy Unit at NUS Yong Loo Lin School of Medicine for imaging training and assistance. We are grateful to Professor Paul Elkington (University of Southampton, UK) for commenting on the manuscript.

## Contributions

C.W.M.O. conceived, obtained funding and analyzed the data. X.Y.P., F.K.L., C.B., H.T.C., J.M.H., Q.H.M., P.M.T. and Y.W. conducted the experiments. S.C.T. and T.C.C.L. performed neuroradiology analysis. S.M.F. screened and recruited the pediatric patients. M.K. and T.W.Y. collected clinical data and biological samples. C.L.T., C.S.L.D. and M.L. provided and reviewed the human CNS-TB brain biopsy specimens. W.K.T., F.K.L. and J.K.T. contributed to LCM RNA-Seq experiment and analysis. F.K.L., A.F.V., J.K.T. and W.K.T. performed bioinformatics analysis of spatial transcriptomics data. R.R. evaluated the histopathology of mouse tissues. L.S.P. and C.L.D. contributed to LC-MS/MS assay development, experimental design, sample analysis and quantitation of TB drugs. W.D.L., E.S.H.C. and G.K.B. contributed to MALDI-MSI experiment and analysis. F.K.L., H.T.C. and T.H.H. analyzed the number of granulomas. F.K.L., B.V., S.C. and Y.F.P. contributed to vasculature analysis. X.Y.P., F.K.L. and C.W.M.O. wrote the first draft of the paper which was reviewed and critically revised by all authors.

## Ethics declarations

Competing interests

CWMO received speaking fees from Qiagen and conference sponsorship from MSD outside this work. All other authors declared no competing interests.

---

**Supplementary Fig. 1.**
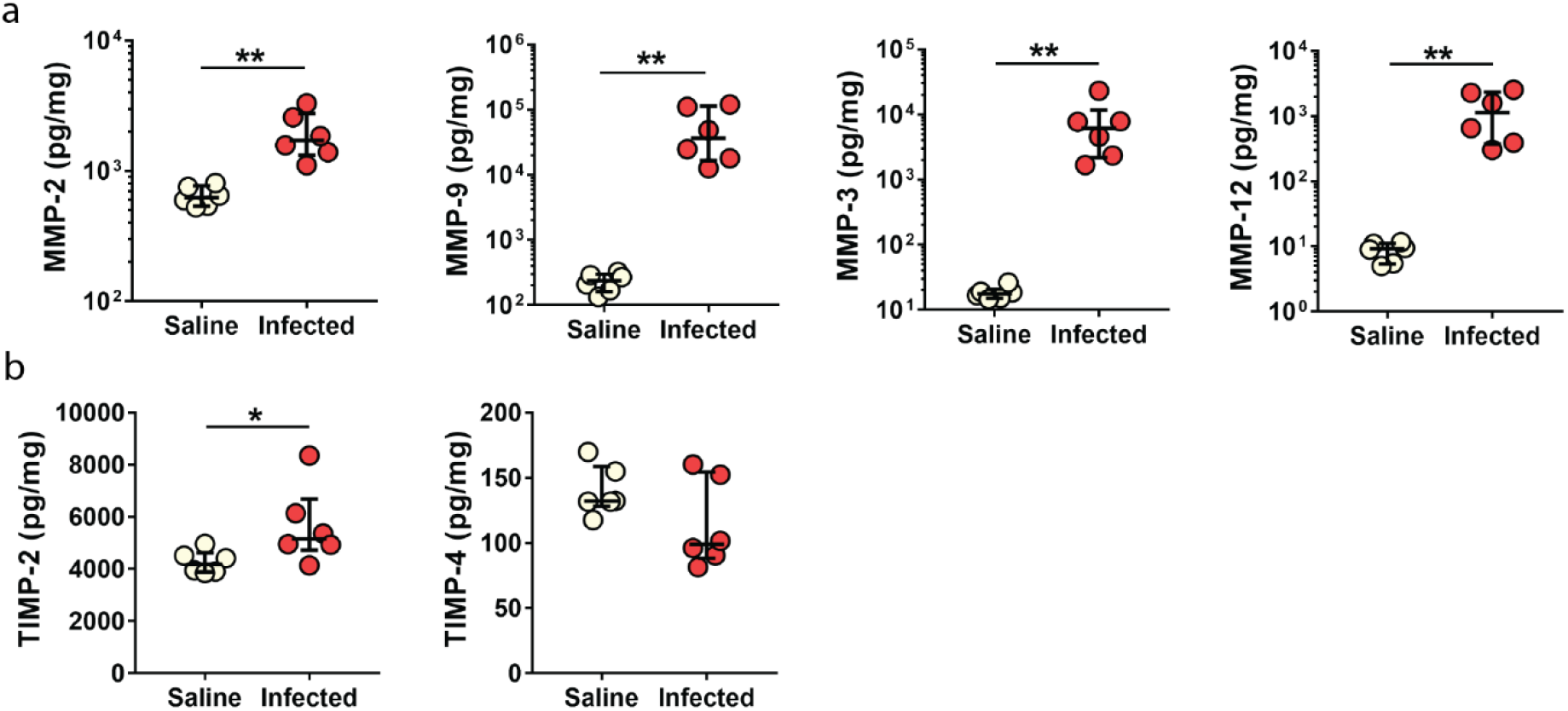
Brain concentrations of (**a**) gelatinase MMP-2, -9, stromelysin MMP-3, and elastase MMP-12, and (**b**) TIMP-2 and -4 in saline control versus H37Rv i.c.vent.-infected *Nos2^-/-^* mice. MMP and TIMP concentrations are normalized to total protein concentration. Bars represent median ± IQR, and analysis was conducted using Mann-Whitney test.

**Supplementary Fig. 2.**
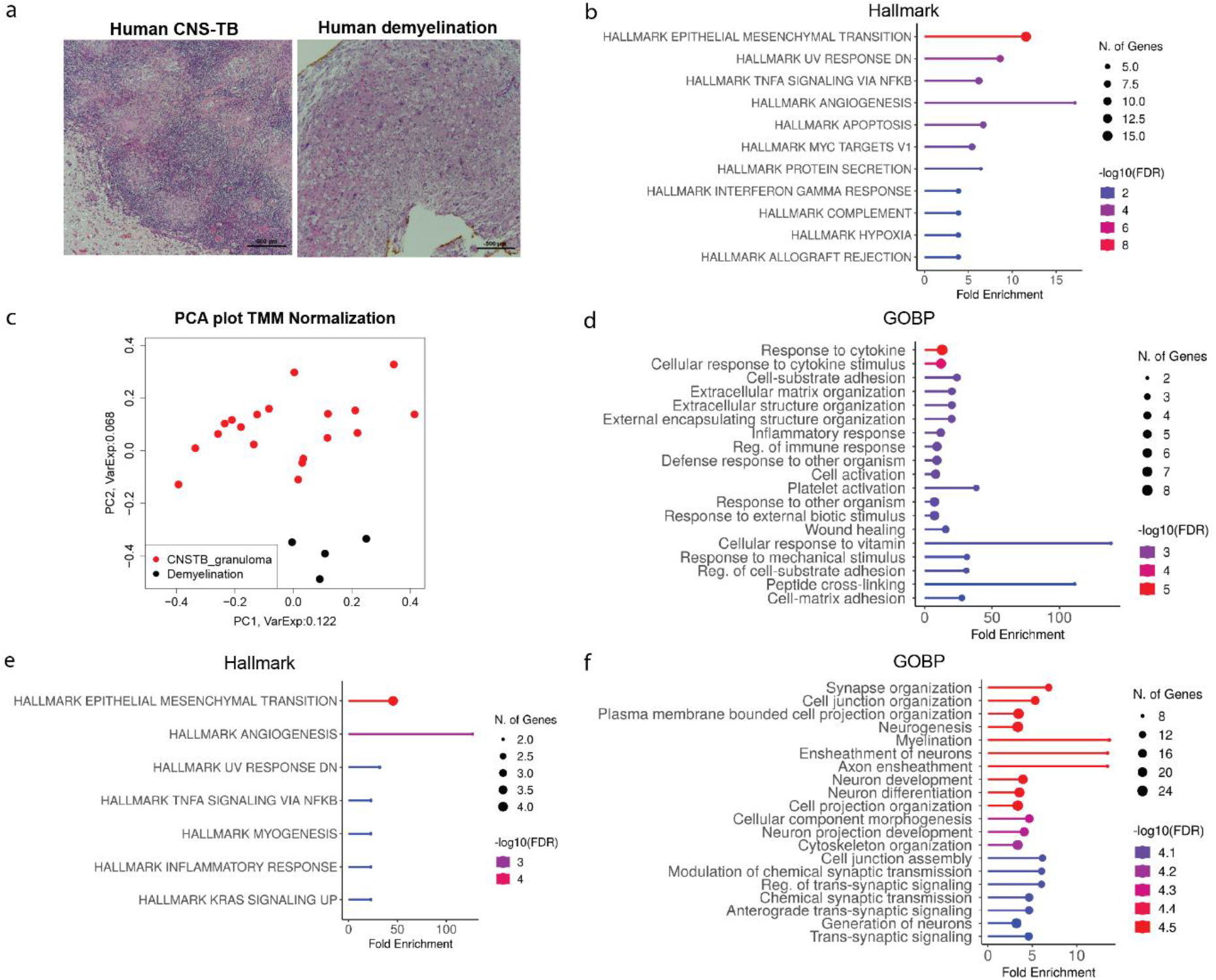
**a**, Hematoxylin-eosin stains of a representative human CNS-TB granuloma (left panel) and human demyelination tissue (right panel). Scale bar = 500 µm. **b**, Top hallmark processes associate with distinct gene cluster comparing human CNS-TB granulomas with other tissue types (demyelination, CNS-TB normal, CNS-TB inflammatory). **c**, PCA plot of gene expression libraries demonstrates clear separation of human demyelination tissue from human CNS-TB granulomas. **d**, Top biological processes enriched in human CNS-TB granulomas compared to human demyelination tissues. **e**, Top hallmark processes enriched in human CNS-TB granulomas compared to human demyelination tissues. **f**, Top biological processes enriched in human demyelination tissues compared to human CNS-TB granulomas. All pathways in this figure were generated using ShinyGO v0.77.

**Supplementary Fig. 3.**
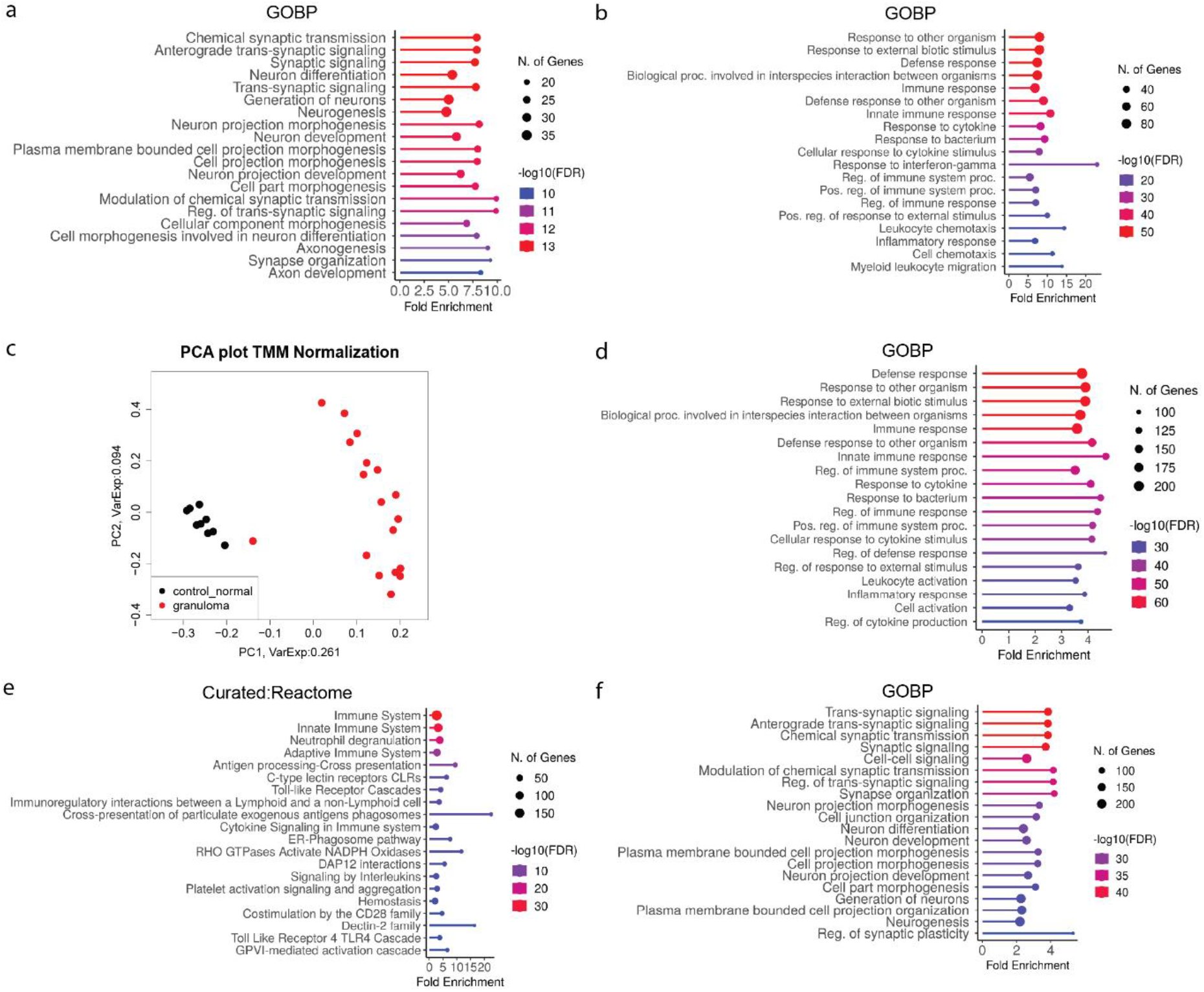
Top biological processes associated with (**a**) down-regulated gene cluster and (**b**) up-regulated gene cluster in murine CNS-TB granulomas compared to other tissue types (control normal, CNS-TB normal, CNS-TB inflammatory). **c**, PCA plot of gene expression libraries showing distinct clustering of samples from murine CNS-TB granulomas and normal tissue from healthy murine controls. **d,** Top biological processes enriched in murine CNS-TB granulomas compared to normal tissues from healthy murine controls. **e**, Top reactome pathways enriched in murine CNS-TB granulomas compared to normal tissues from healthy murine controls. **f**, Top biological processes enriched in normal tissues from healthy murine controls compared to murine CNS-TB granulomas. All pathways in this figure were generated using ShinyGO v0.77.

**Supplementary Fig. 4.**
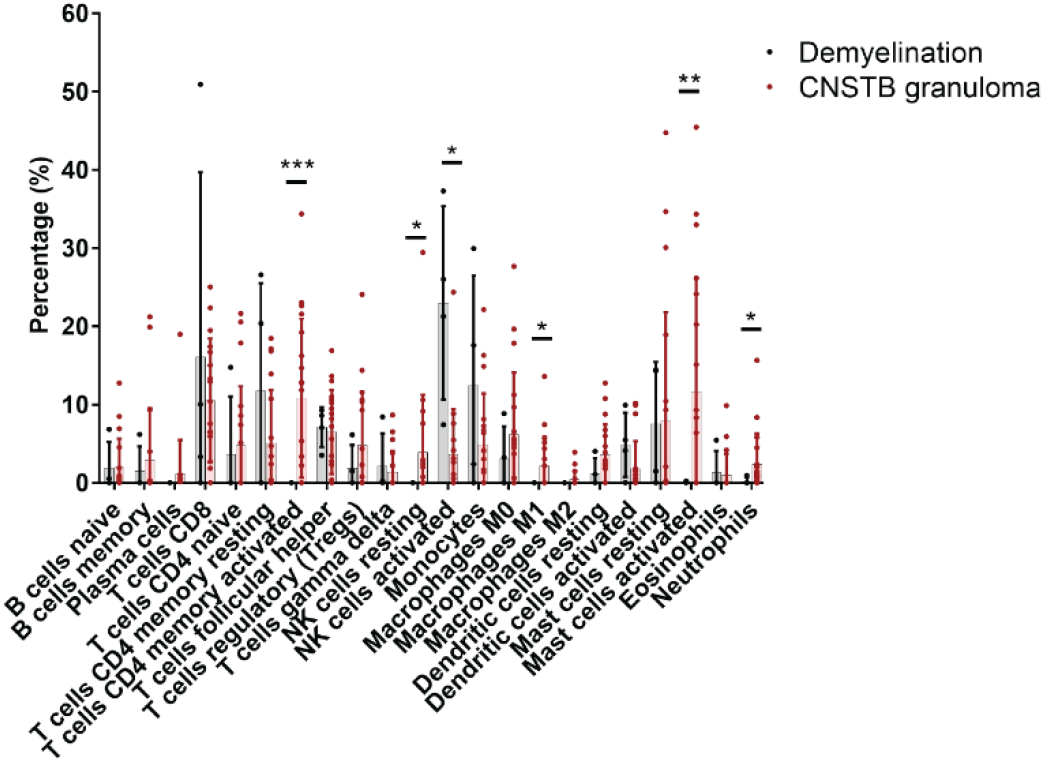
Deconvolution analysis using LM22 signature matrix was performed to compare the cell fractions in human demyelination tissues and human CNS-TB granulomas. Bars represent mean ± SD, and analysis was conducted using unpaired t test with Welch’s correction. * *P* < 0.05; ** *P* < 0.01; *** *P* < 0.001; **** *P* < 0.0001.

**Supplementary Fig. 5.**
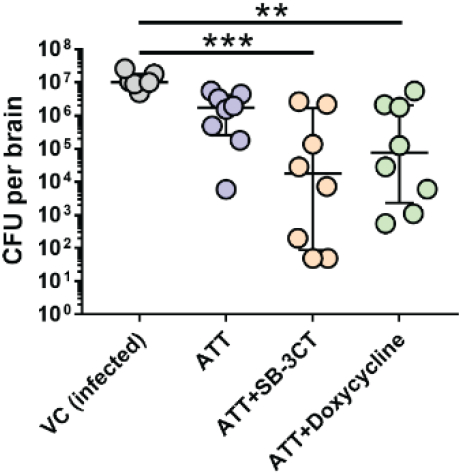
CFU enumeration in brain homogenates shows significantly reduced *M. tb* burden in ATT+SB-3CT and ATT+Doxycycline groups. Bars represent median ± IQR. Analysis by Kruskal-Wallis test with Dunn’s multiple comparisons test. ** *P* < 0.01; *** *P* < 0.001.

**Supplementary Fig. 6.**
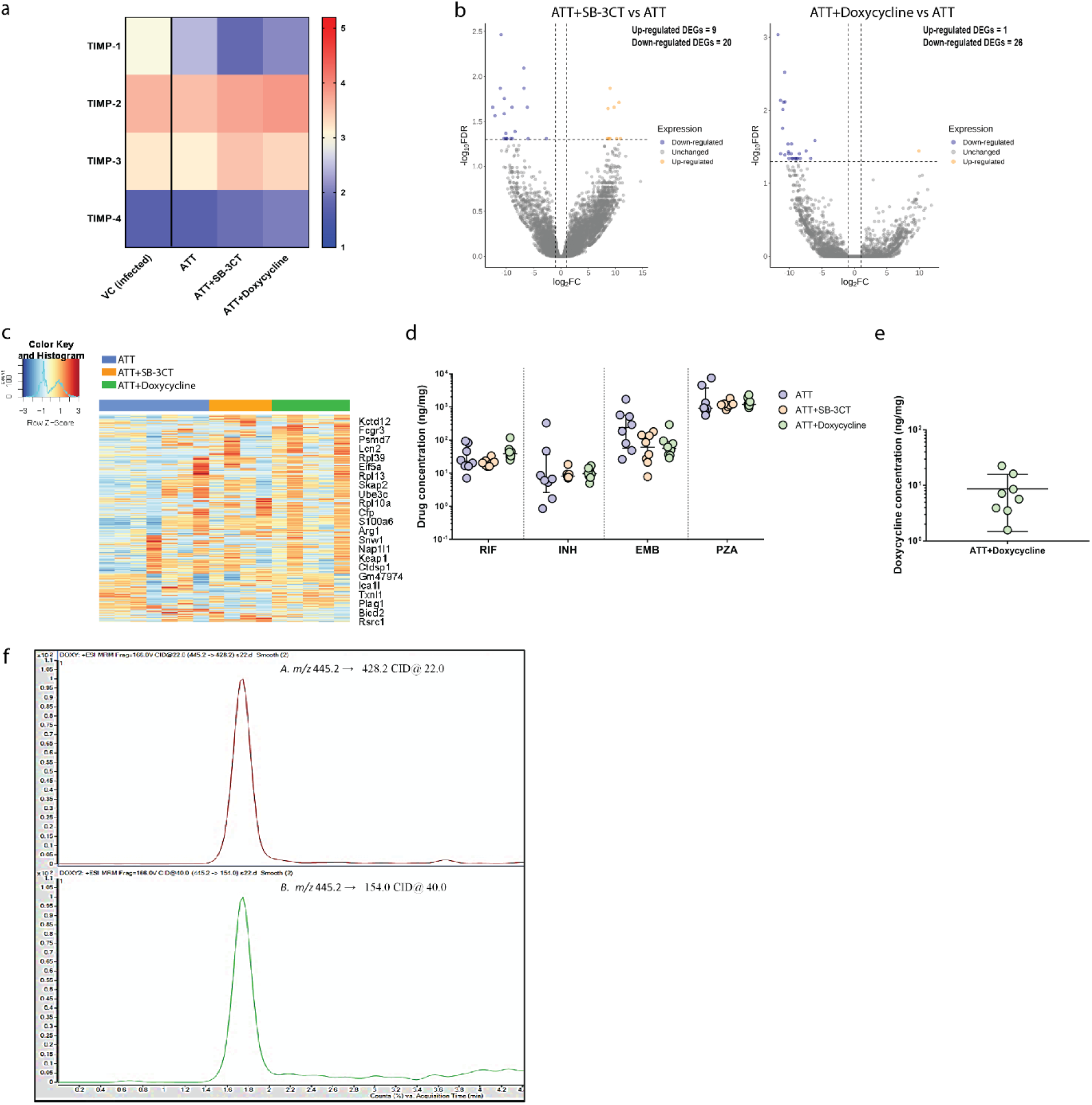
**a**, Heatmap showing median expression of TIMP-1, -2, -3 and -4 in the brain homogenates of CNS-TB mice from different treatment groups. TIMP concentrations are normalized to total protein concentration. **b**, Volcano plot shows up-regulated (log fold-change >2, *FDR*<0.05) and down-regulated (log fold-change <-2, *FDR* <0.05) genes of CNS-TB mice treated with ATT+SB-3CT versus ATT alone (left panel). Volcano plot shows up-regulated (log fold-change >2, *FDR*<0.05) and down-regulated (log fold-change <-2, *FDR* <0.05) genes of murine CNS-TB that received ATT+Doxycycline in comparison with ATT alone (right panel). **c,** Hierarchical clustering heatmap of top 500 most variable genes using Pearson correlation and Ward method. The gene expressions are heterogeneous among the three murine CNS-TB groups. **d,** Rifampicin, Isoniazid, Ethambutol and Pyrazinamide are detected in the brain homogenates of *M. tb*-infected mice with comparable concentrations across the groups. Bars represent median ± IQR and analysis was conducted using Kruskal-Wallis test with Dunn’s multiple comparisons test. **e**, Concentration of doxycycline was measured in the brain homogenates of *M. tb*-infected mice treated with ATT and doxycycline. Bars represent mean ± SD, analysis was conducted using one-way ANOVA with Dunn’s multiple comparisons test. **f,** Extracted MRM chromatograms of doxycycline quantifier (top panel) and qualifier (bottom panel). The X-axis denotes the retention time, tR (min), and the Y-axis denotes ion intensity, from which the area under the curve (AUC) was obtained by integration.

**Supplementary Table 1.** Infectious etiologies in the human pediatric cohort.

|  | Number |
| --- | --- |
| **Viral meningitis** | **n = 10** |
| Japanese Encephalitis virus | 3 |
| Adenovirus | 1 |
| Clinically-diagnosed | 6 |
| **Bacterial meningitis** | **n = 24** |
| *Haemophilus Influenzae Type B (Hib)* | 8 |
| *Streptococcus pneumoniae (S. pneumoniae)* | 3 |
| Hib + *S. pneumoniae* | 1 |
| *Staphylococcus aureus* | 2 |
| *Salmonella typhi* | 1 |
| *Escherichia coli* | 1 |
| *Klebsiella pneumoniae* | 1 |
| *Neisseria meningitidis* | 2 |
| *Bordetella pertussis* | 1 |
| Mycoplasma | 1 |
| Clinically-diagnosed | 3 |
| **Tuberculous meningitis** | **n = 20** |
| Definite | 2 |
| Clinically diagnosed and treated | 18 |

**Supplementary Table 2.**
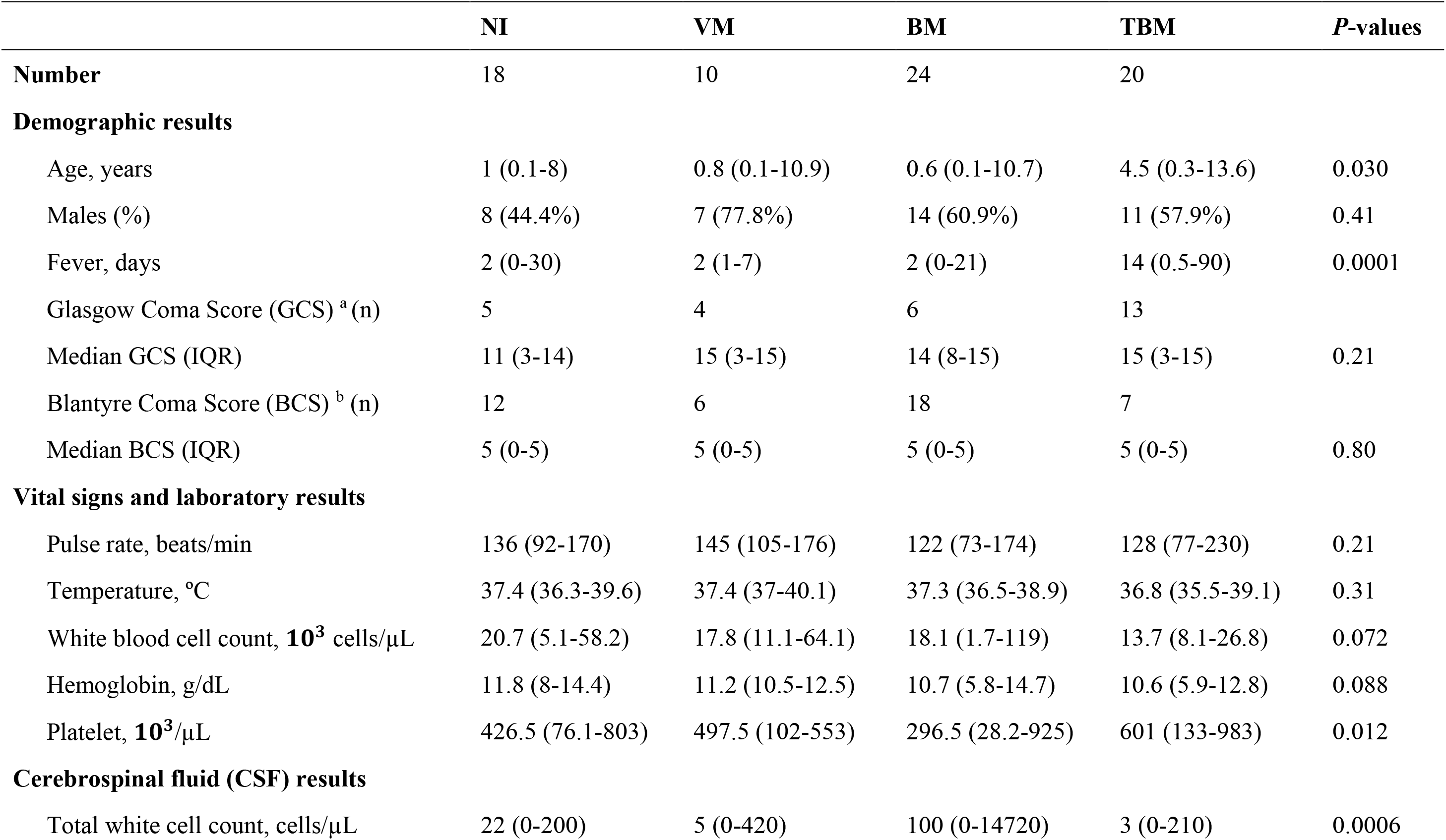
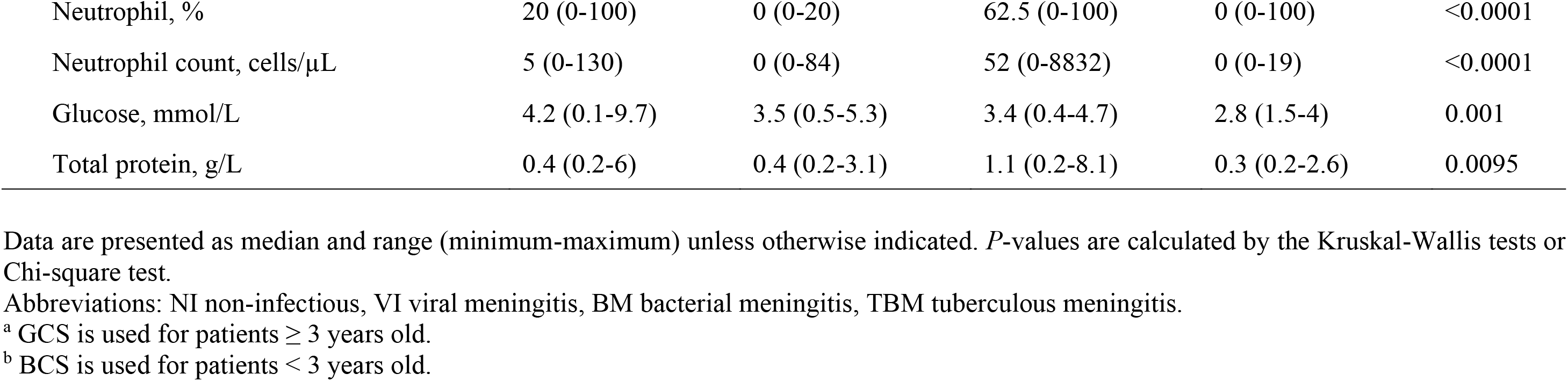
Baseline clinical data and cerebrospinal fluid results of the human pediatric cohort.

**Supplementary Table 3.** Histopathological evaluation of *M. tb*- lesions in H37Rv-infected *Nos2^−/−^* mice from different treatment groups, showing smaller scores indicative of less immunopathology in adjunctively treated SB-3CT and doxycycline groups.

| Lesions <sup>a</sup> | VC (infected) |  |  |  |  | ATT |  |  |  |  | ATT+SB-3CT |  |  |  |  | ATT+Doxycycline |  |  |  |  |
| --- | --- | --- | --- | --- | --- | --- | --- | --- | --- | --- | --- | --- | --- | --- | --- | --- | --- | --- | --- | --- |
|  | M | C | H | T | n | M | C | H | T | n | M | C | H | T | n | M | C | H | T | n |
| Inflammation (MNCs)/PMNCs | 1 | 1 | 1 | 0 | **6** | 1 | 1 | 1 | 0 | **8** | 0 | 1 | 1 | 0 | **3** | 1 | 0 | 1 | 0 | **4** |
| Perivascular cuffing | 0 | 1 | 1 | 0 | **6** | 1 | 2 | 1 | 0 | **8** | 0 | 0 | 1 | 0 | **4** | 0 | 1 | 1 | 0 | **4** |
| Gliosis | 0 | 1 | 1 | 0 | **6** | NA | 1 | 0 | 0 | **4** | NA | 0 | 0 | 0 | **1** | NA | 0 | 0 | 0 | **1** |
| Granuloma (not well formed) | 0 | 0 | 0 | 0 | **1** | 0 | 0 | 0 | 0 | **1** | 0 | 0 | 1 | 0 | **3** | 0 | 0 | 1 | 0 | **3** |
| Pyogranuloma | 0 | 1 | 1 | 0 | **6** | 0 | 1 | 1 | 0 | **5** | 0 | 0 | 0 | 0 | **0** | 0 | 0 | 0 | 0 | **1** |
| Neuronal degeneration/necrosis | 0 | 0 | 0 | 0 | **4** | NA | 0 | 0 | 0 | **3** | NA | 0 | 0 | 0 | **0** | NA | 0 | 0 | 0 | **2** |
| Liquefactive necrosis (±) <sup>b</sup> | - | + | + | - | **3** | - | + | + | - | **3** | - | - | - | - | **0** | - | - | - | - | **0** |
| Presence of bacilli (±) <sup>b</sup> | - | + | + | + | **6** | - | + | + | - | **5** | - | - | + | - | **1** | - | - | + | - | **1** |

Abbreviations: *M* meninges, *C* cerebral cortex, *H* hippocampus, *T* thalamus, *n* number of mice present the lesions, *MNCs* mononuclear cells, *PMNCs* polymorphonuclear cells, *NA* not applicable
<sup>a</sup> Severity of lesions in each group are scored on a scale of 0–5: 0 – no abnormalities detected; 1 – minimal; 2 – mild; 3 – moderate; 4 – marked; 5 – severe. The average score of 7 to 8 mice per group is shown.
<sup>b</sup> + represents present, – represents absent

**Supplementary Table 4.** Multiple reaction monitoring transitions of drugs tested using LC-MS/MS.

| **Compound** | **Average**<br>**Molecular**<br>**Mass** | **MRM**<br>**Transition**<br>**(m/z)** | **Retention**<br>**Time (min)** | **Collision**<br>**Energy (eV)** |
| --- | --- | --- | --- | --- |
| Rifampicin | 822.9 | 823.4 > 791.3 | 2.695 | 16 |
|  |  | 823.4 > 163.1 | 2.695 | 41 |
| Doxycycline | 444.4 | 445.2 > 428.8 | 1.749 | 22 |
|  |  | 445.2 > 154.0 | 1.749 | 40 |
| Ethambutol | 204.3 | 205.1 > 116.1 | 0.543 | 8 |
| Isoniazid | 137.1 | 138.1 > 121.1 | 0.671 | 8 |
| Pyrazinamide | 123.1 | 124.1 > 81.1 | 0.922 | 17 |
| SB-3CT | 306.4 | 307.1 > 232.8 | 3.388 | 5 |

**Supplementary File 1** List of genes for heatmap clusters of top 500 variable genes from human samples

**Supplementary File 2** List of significant DEGs for human CNS-TB granulomas versus human demyelination samples

**Supplementary File 3** List of genes for heatmap clusters of top 500 variable genes from murine samples

**Supplementary File 4** List of significant DEGs for murine CNS-TB granulomas versus normal tissues from healthy murine

**Supplementary File 5** List of neutrophil genes for heatmap clusters from murine healthy brain tissues versus murine CNS-TB granulomas

**Supplementary File 6** List of genes for heatmap clusters of top 500 variable genes from murine CNS-TB received different treatment regimes

## Notes

### Competing Interest Statement

The authors have declared no competing interest.

